# Metabolic connectivity in ageing

**DOI:** 10.1101/2024.06.16.599247

**Authors:** Hamish A. Deery, Emma Liang, M. Navyaan Siddiqui, Gerard Murray, Katharina Voigt, Robert Di Paolo, Chris Moran, Gary F. Egan, Sharna D. Jamadar

## Abstract

Information transfer across the brain has a high energetic cost and requires the efficient use of glucose. Positron emission tomography (PET) studies have shown that ageing is associated with a decline in regional rates of cerebral glucose metabolism. However, until recently, it has not been possible to measure the timecourse of molecular activity within an individual using PET, preventing the study of metabolic network connectivity across the brain. Here we report the results of the first high temporal resolution functional PET study examining metabolic connectivity and cognitive function in ageing. The metabolic connectomes of 40 younger (mean age 27.9 years; range 20-42) and 46 older (mean 75.8; 60-89) adults were characterised by high connectivity strength in the frontal, temporal, motor, parietal and medial cortices. Ageing was associated with lower global integration of metabolic hub regions, indicating disrupted information transfer across the metabolic network in older adults. In younger adults, a high proportion of glucose was used to support hubs in the frontal regions. Older adults had a smaller energy budget in comparison to younger adults, and older adults used a higher proportion of energy to support mostly posterior hub regions. This difference in the metabolic network topology in older adults was associated with worse cognitive performance. We conclude that ageing is associated with reduced metabolic connectivity, an altered metabolic network topology and a high glucose cost in hub regions. Our results highlight the fundamental role that metabolism plays in supporting information transfer in the brain and the unique insights that metabolic connectivity provides into the ageing brain.

## Introduction

The human brain relies on a large energy budget [1]. The brain accounts for only 2% of the body’s total weight but 20% of whole body resting metabolism [2, 3]. The majority of this energy is generated through the oxidative metabolism of glucose [4]. A significant proportion of glucose metabolism in the brain is used to maintain the functional resting state [5]. Age-related decline in cognitive function and neurodegeneration [6] have been associated with decline in cerebral metabolism [7] and have led to interest in how glucodynamics and the coherence of metabolism in brain networks change during ageing [5].

Understanding brain glucose metabolism requires knowledge of the underlying neural activity and network connectivity of the brain’s resting state. To date, most work on the neural bases of resting state network connectivity has largely used blood oxygenation level dependent functional magnetic resonance imaging (BOLD-fMRI or fMRI). From a brain energetics perspective, the utility of fMRI stems from its ability to index neurovascular coupling [8]. A limitation of fMRI is that it does not directly measure the metabolic costs of neural activity or maintenance of the resting state. As such, previous fMRI connectome research has shed little light on the metabolic component of neurovascular coupling. [18F]-fluorodeoxyglucose positron emission tomography (FDG-PET), and its more recent variant FDG-functional PET (FDG-fPET, or fPET), can provide a direct and quantitative measurement of the metabolic costs of brain connectivity *in vivo*. To date, age differences in brain connectivity using PET have been based on across-subject covariance of regional cerebral metabolic rates of glucose (CMR_GLC_) rather than within-subject timeseries correlation as is standard in fMRI and EEG [9]^1^. These studies have reported that older people display lower metabolic covariance within networks, and a more heterogenous pattern of high and low correlations between-networks compared to younger adults [10-13].

However, group-level differences in CMR_GLC_ covariance cannot index information transfer within the brain network [14-17]. Furthermore, they do not adequately reflect within-subject metabolic connectivity [17, 18]. Estimation of the metabolic connectivity of an individual person requires a timeseries of glucodynamics, such as that provided by fPET [19-22]. Previous work in younger adults using fPET has reported findings different to those using fMRI, particularly their associations with executive function [23]. In younger adults, metabolic connectomes derived from fPET have shown strong frontoparietal connections both within and between hemispheres [17]. When compared to fMRI derived connectomes, fPET connectomes are uniquely related to executive function [23]. However, to date, metabolic connectomes have not been studied in older adults.

A organisational principle that governs the human connectome is a trade-off between minimisation of the network energy cost and maximisation of communication efficiency [24]. Wiring cost refers to the physiological resource or substrates that are needed to support network connections. The longer and denser the connections between nodes in a network, the higher the wiring cost of the network [25]. Activity cost is the energy consumed in neural signal activity [26]. Cost minimisation principles constrain the possible number, distance and strength of connections in a brain network to local regional nodes more likely to connect to each other, rather than regions requiring longer distance connections. This results in high connectivity strength between neighbouring nodes, leading to networks that are characterised by local clustering, local efficiency and network segregation [27, 28]. However, cost minimisation principles are insufficient to describe the human brain network. Long-distance connections also play an important role in integration of distributed functional networks [24, 29, 30]. As a result, there is a trade-off between cost minimisation and other network properties that facilitate system-wide function, such as global integration and efficiency. Based on fMRI, the networks of older adults are less segregated and efficient but more integrated than the networks of younger adults [see [31] for review].

Metabolic hubs are likely to be of particular importance for optimal brain function because they are highly connected via long-distance neural pathways to support information integration and coordination [24, 29, 32]. Hubs of functional brain networks are initially located in primary processing regions in early childhood, and in association areas by late childhood [33]. Hubs have been reported in the frontal and parietal cortices, hippocampus, precuneus and subcortical structures [32, 33]. Hubs have high rates of cerebral blood flow and aerobic glycolysis, suggesting that they have a high energy requirement, although with associated high value for the integration of information processing [29]. Using structural and functional MRI, it has been shown that older adults display reduced hub connectivity strength, degree and centrality, particularly in regions in the frontal and parietal hubs compared with younger adults, indicating a reduced capacity of information transfer to and from the hubs and other regions [30, 34-37].

Here, we use fPET to construct and analyse the metabolic connectome, investigate its alterations from younger to older adulthood, and its relationship to cognition. We measure the cost of metabolic networks using graph metrics and regional rates of glucose metabolism, that mechanistically reflect the cost of glucose metabolism in synaptic activity [38]. We hypothesise that differences between younger and older adults in metabolic connectivity will be observed predominantly in frontal and temporal regions, consistent with earlier studies of cerebral glucose metabolism in ageing [7]. Secondly, metabolic connectomes will show lower within- and higher between region network connectivity among older than younger adults [31]. Thirdly, that the metabolic hubs of older adults will show lower efficiency, degree and centrality than the hubs of younger adults [31]. Fourthly, we hypothesise that the rate of glucose metabolism will be higher in hub than non-hub regions, and that a greater proportion of brain glucose metabolism will support network connections in hubs than non-hubs and to a greater extent in older than younger adults. Finally, we hypothesise that higher efficiency, degree and centrality of metabolic hubs will be associated with better cognitive performance and faster processing speed.

## Results

### Sample characteristics

The characteristics of the younger (N = 40) and older (N = 46) participants are shown in Table 1. The mean age of the younger group was 27.9 years (SD=6.2, range 20 – 42 years) and the older group 75.8 years (SD=6.0, range 60-89 years). The proportion of women was higher in the younger group (54%) than the older group (48%) but this difference was not statistically significant (p=.662). The average BMI was similar in the younger (24.1 kg/m^2^) and older (25.7kg/m^2^) groups. The average years of education was higher in the younger group (18.0) than the older group (16.3). Mean fasting blood glucose was greater in the older (5.2 mmol/L) than the younger (4.8 mmol/L) group (p<.001).

**Table 1.**
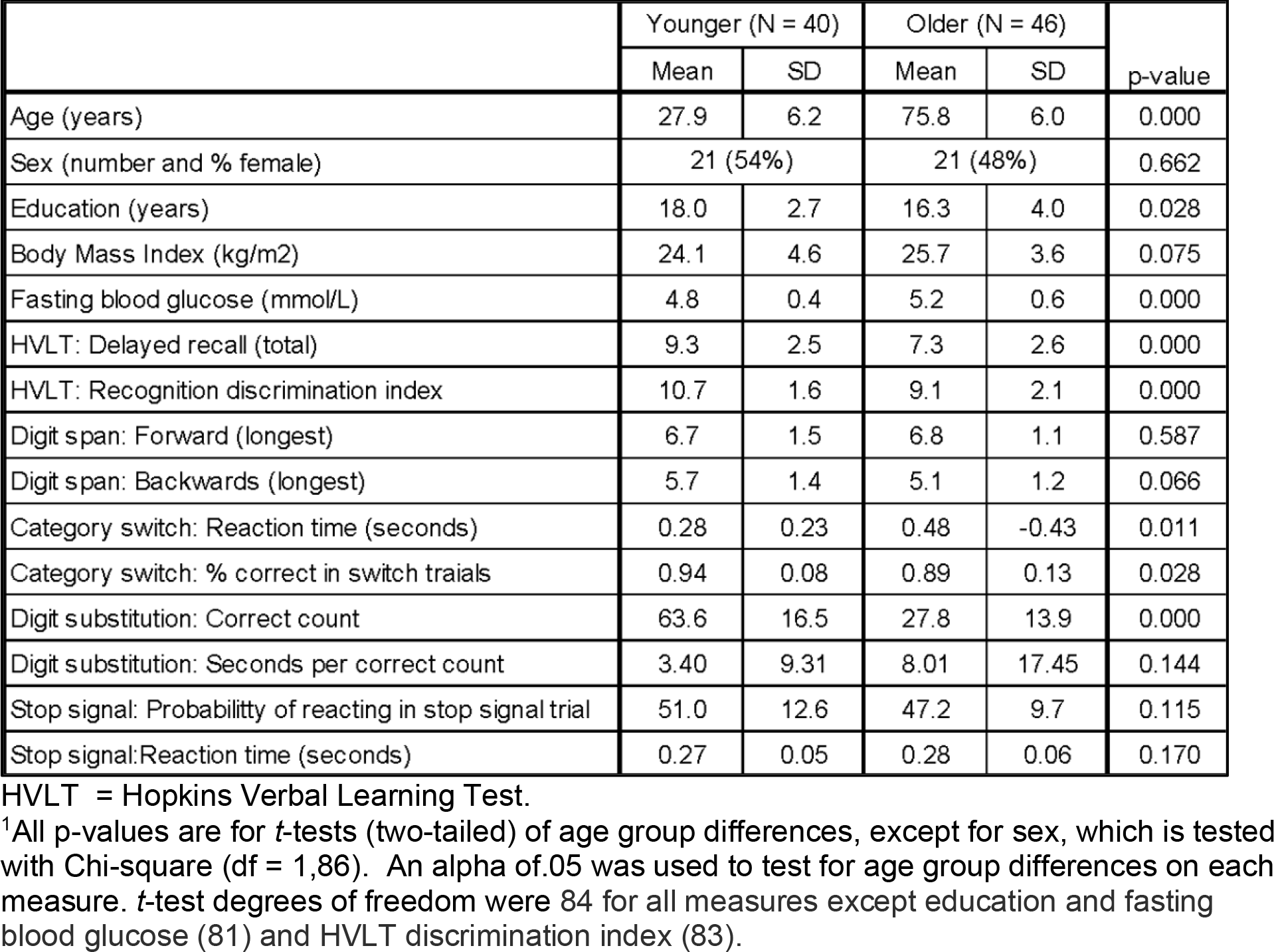
Mean and standard deviation (SD) of demographic and cognitive measures for older and younger adults, and p-value of statistical tests of age group differences^1^.

|  | Younger (N = 40) |  | Older (N = 46) |  | p-value |
| --- | --- | --- | --- | --- | --- |
|  | Mean | SD | Mean | SD |  |
| Age (years) | 27.9 | 6.2 | 75.8 | 6.0 | 0.000 |
| Sex (number and % female) | 21 (54%) |  | 21 (48%) |  | 0.662 |
| Education (years) | 18.0 | 2.7 | 16.3 | 4.0 | 0.028 |
| Body Mass Index (kg/m<sup>2</sup>) | 24.1 | 4.6 | 25.7 | 3.6 | 0.075 |
| Fasting blood glucose (mmol/L) | 4.8 | 0.4 | 5.2 | 0.6 | 0.000 |
| HVLT: Delayed recall (total) | 9.3 | 2.5 | 7.3 | 2.6 | 0.000 |
| HVLT: Recognition discrimination index | 10.7 | 1.6 | 9.1 | 2.1 | 0.000 |
| Digit span: Forward (longest) | 6.7 | 1.5 | 6.8 | 1.1 | 0.587 |
| Digit span: Backwards (longest) | 5.7 | 1.4 | 5.1 | 1.2 | 0.066 |
| Category switch: Reaction time (seconds) | 0.28 | 0.23 | 0.48 | -0.43 | 0.011 |
| Category switch: % correct in switch trials | 0.94 | 0.08 | 0.89 | 0.13 | 0.028 |
| Digit substitution: Correct count | 63.6 | 16.5 | 27.8 | 13.9 | 0.000 |
| Digit substitution: Seconds per correct count | 3.40 | 9.31 | 8.01 | 17.45 | 0.144 |
| Stop signal: Probability of reacting in stop signal trial | 51.0 | 12.6 | 47.2 | 9.7 | 0.115 |
| Stop signal: Reaction time (seconds) | 0.27 | 0.05 | 0.28 | 0.06 | 0.170 |

HVLT = Hopkins Verbal Learning Test.
<sup>1</sup>All p-values are for *t*-tests (two-tailed) of age group differences, except for sex, which is tested with Chi-square (df = 1,86). An alpha of .05 was used to test for age group differences on each measure. *t*-test degrees of freedom were 84 for all measures except education and fasting blood glucose (81) and HVLT discrimination index (83).

Overall, the older group showed poorer memory, processing speed and cognitive flexibility compared to the younger group. In particular, the older group showed lower delayed recall and recognition discrimination index in the verbal learning test than did younger adults (both p < .001). Older adults also had a lower percentage of correct switch trials than younger adults (p < .05). Compared to younger adults, older adults had lower correct count (p < .001) in the digit substitution task and a slower reaction time in the category switch task (p < .05). There were no significant age differences in performance on the digit span and stop signal tasks.

### Age Differences in Metabolic Connectomes

Metabolic connectomes were derived by cross-correlating the timeseries of fPET signal between 106 Harvard-Oxford atlas regions (see Methods). An anatomical parcellation was chosen to define the metabolic connectome, as it is unclear if ‘functional’ atlases derived from fMRI data are applicable to metabolic data [17, 19, 20, 39]. In the Supplement, we present results for the Schaefer 100 fMRI-derived atlas, which indicates a compatible pattern of effects to those presented here.

The most salient qualitative characteristic for the group-averaged metabolic connectome for both younger and older adults was high connectivity strength for the regions in frontal, motor and parietal cortices, both within and between hemispheres (Figure 1A and 1B). Medial connectivity between the frontal, motor and parietal regions was also evident for the younger and older adults. In contrast, between connectivity strength was relatively low for the limbic, temporal, lateral temporal cortices and subcortex for both younger and older adults. A second salient feature is the apparent lower connectivity strength across the connectome for older than younger adults, as seen by less dark red-to-orange, particularly for the frontal, medial and parietal cortices.

**Figure 1.**
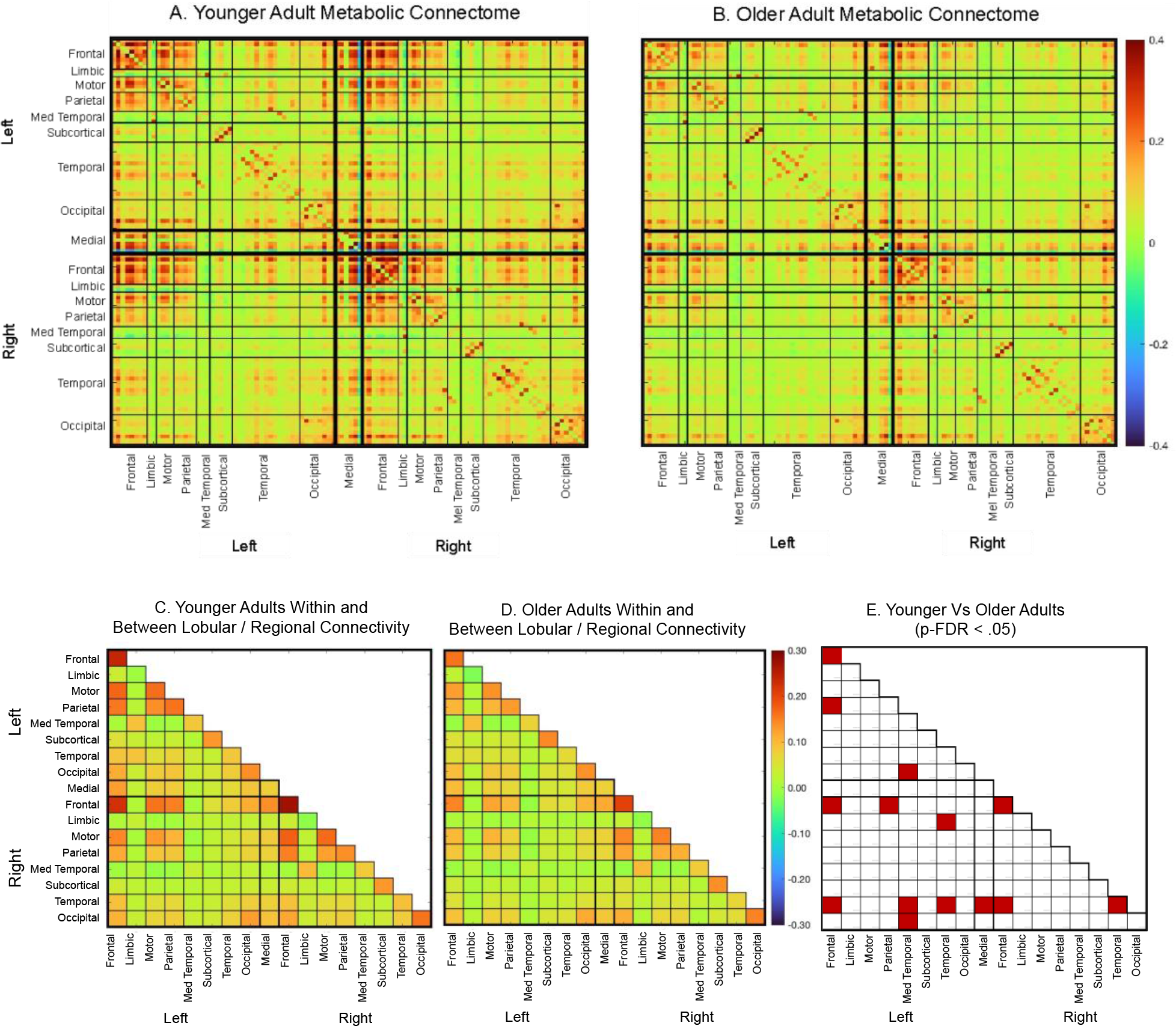
Metabolic connectivity for younger and older adults. (A) Younger and (B) older adult metabolic connectomes across the 106 nodes in the network. Within and between lobular / regional connectivity averaging the nodes within the lobes for (C) younger and (D) older adults. Within connectivity is shown in the diagonal cells and between connectivity on the off-diagonal cells of each matrix. (E) significance test (*t*-test, df = 84) of younger vs older adults, with red shaded cells indicating a statistically significant age group differences at p-FDR<.05. All age group differences are lower connectivity for older vs younger adults..

### Age Differences in Within and Between Region Connectivity

The topology of connections noted above for younger and older adults was evident when within- and between connectivity was calculated (Figures 1C and 1D). Compared to younger adults, older adults showed significantly lower within connectivity among the regions in the left and right frontal cortices and the right temporal cortex (Figure 1D). The left and right frontal cortices showed lower connectivity between their respective left and right parietal regions in older than younger adults. Lower connectivity between the left occipital regions and the right medial temporal regions was seen in older adults. Older adults also had lower connectivity between the right temporal lobe and the left medial temporal and left temporal regions, as well as the right frontal lobe regions. Older adults also showed lower connectivity between the right limbic and left temporal regiona than younger adults.

### Metabolic Cost and Age Differences in the Function of Hub Regions

Since fMRI studies have shown that the “wiring cost” of functional connectivity increases monotonically with increasing connection density [24], we examined the topology of the metabolic connectome at two levels of network degree. At the top 10% of network edges, thirteen regions were identified as metabolic hubs in the frontal, motor, occipital and medial cortices (see Supplementary Information, Figure S2). The regions identified as metabolic hubs included the left and right frontal poles, bilateral middle frontal gyri and right superior frontal gyrus in the frontal lobes; the left and right precentral gyri and left postcentral gyri in the motor cortex; the bilateral superior division of the lateral occipital cortices and the right occipital pole in the occipital lobe; and the posterior division of the cingulate gyrus and precuneus in the medial cortex. At the top 30% of network edges, two additional regions were identified as hubs: the right frontal gyrus and frontal orbital cortex (Figure S3).

Significant age group differences were found in the multivariate test comparing the hubs of younger and older adults on global efficiency (*F*(15,70) = 2.0, p < .05), local efficiency (*F*15,70) = 3.4, p = .001), betweenness centrality (*F*(15,70) = 2.9, p = .002) and degree (*F*(15,70) = 2.7, p = .003) at the top 10% of edges. Age differences were also found at the top 30% of edges for global efficiency (*F*(15,70) = 13.6, p < .001), local efficiency (*F*(15,70) = 4.4, p < .001), betweenness centrality, (*F*(15,70) = 3.5, p < .001) and degree (*F*(15,70) = 3.0, p = .001). The follow-up univariate tests at the top 10% of edges revealed that older adults had higher global efficiency in hub regions in the motor, occipital, and medial cortices, namely, the left postcentral gyrus and lateral occipital cortex, the right occipital pole and the precuneus and cingulate gyri (Figure 2A.i and Table S1). However, at the top 30% of edges, the direction of the age differences reversed (Figure 2B.i). Younger adults showed higher global efficiency than older adults in the left and right frontal poles and right superior frontal and middle frontal gyri. The age differences in betweenness centrality largely followed a similar pattern to global efficiency (Figure 2A.iii and 2B.iii). For degree, the age group differences were in the frontal hub regions at both the top 10% (Figure 2A.iv) and top 30% (Figure 2B.iv) of edges. Older adults showed lower degree than younger adults in the bilateral frontal poles and middle frontal gyri, and the right superior frontal gyrus.

**Figure 2.**
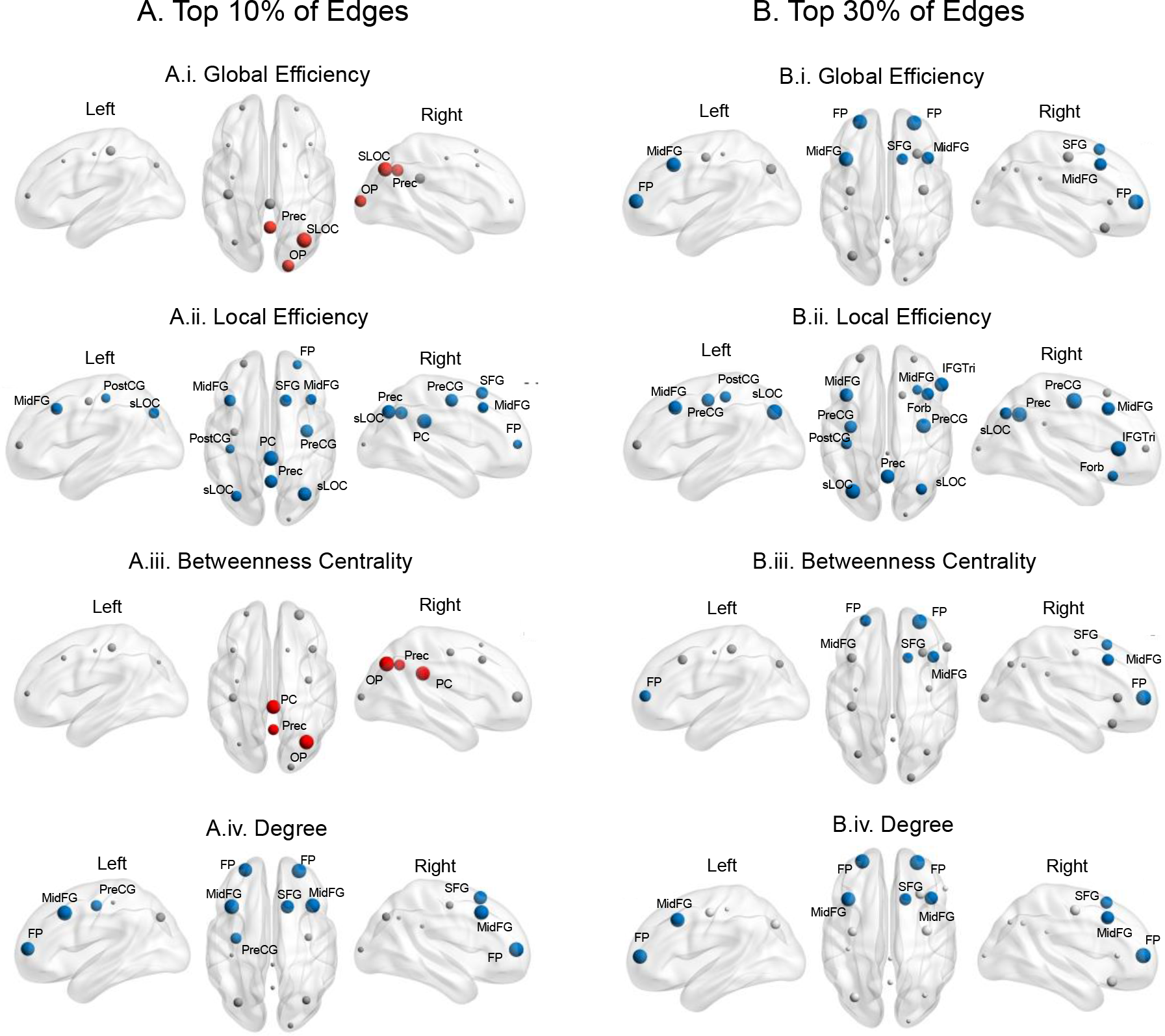
Graph metrics results for *hubs* of the metabolic network comparing younger and older adults at the top 10% (A) and 30% (B) of network edges. Nodes showing significant age group differences are colored red for older > younger and colored blue for younger > older. Nodes showing non-significant age group differences are shaded grey. Relative size of the node reflects the magnitude of the F-statistic (see Table S1). FP = Frontal Pole; MidFG = Middle Frontal Gyrus; SFG = Superior Frontal Gyrus; IFGtri = Inferior Frontal Gyrus; Pars Triangularis; Forb = Frontal Orbital Cortex; RPreCG = Precentral Gyrus; PostCG = Postcentral Gyrus; sLOC = Lateral Occipital Cortex, Superior Division; OP =Occipital Pole; PC = Cingulate Gyrus, Posterior Division; and Prec = Precuneus Cortex. Figures produced using Brain Net Viewer (http://www.nitrc.org/projects/bnv/) (Xia et al., 2013).

Younger adults showed higher local efficiency than older adults in the metabolic hub regions across the brain at both the top 10% and 30% of edges (Figures 2A.ii and 2B.ii), including the right frontal pole, left and right middle frontal gyri and the right superior frontal gyrus, the left and right precentral gyri, lateral occipital cortex, and the precuneus and cingulate gyrus.

The topology of the regional age differences across the whole brain largely followed those for the hubs and are reported in the Supplement (Figure S4 and Tables S2 and S3).

### Cerebral Metabolic Rates of Glucose and Glucose Cost Index of Hub and Non-Hub Regions

Across the sample, the average CMR_GLC_ was 4.1% higher in the 13 hubs than the 93 non-hub regions at the top 10% of edges (*t*(80) = 12.7, p < .001; see Figure S5). The average CMR_GLC_ was higher in the hub than non-hub regions for 83 of 86 participants. At the top 30% of edges, the average CMR_GLC_ across the sample was 3.4% higher in the 15 hubs than the 91 non-hub regions (*t*(80) = 10.8, p < .001). CMR_GLC_ was also higher in the hubs than non-hubs for 78 of 86 participants at the top 30% of edges. The hubs in the frontal and medial cortices showed the highest CMR_GLU_ of the hub regions (see Figure S5).

A Glucose Cost Index (GCI) was defined as the ratio between global efficiency, local efficiency, betweenness centrality, degree, and CMR_GLU_ in each hub region. The GCI values reflect the metabolic cost of the network properties measured by the graph metrics. For example, compared to a region with a lower GCI for global efficiency, a region with a higher GCI has shorter average path length relative to its rate of glucose metabolism, indicating that it is using a larger proportion of glucose to support global integration. The average GCI was significantly higher in the hub than non-hub regions, and significantly higher at the top 30% than top 10% of edge for all graph metrics (see Table S4). Age group differences were found in GCI at the top 10% of edges for global efficiency (*F*(1,85) = 2.7, p = .004), local efficiency (*F*(1,85) = 3.4, p = .001), betweenness centrality (*F*(1,85) = 3.2, p = .001) but not degree (*F*(1,85) = 1.5, p = .158). Age differences were also found in the GCI at the top 30% of edges for global efficiency (*F*(1,85) = 2.0, p = .002), local efficiency (*F*(1,85) = 2.7, p = .003), betweenness centrality (*F*(1,85) = 3.5, p < .001) but not degree (*F*(1,85) = 1.4, p = .176). The hubs showing significant age group differences in the post-hoc univariate tests are shown in Figure 3 (also see Figure S6, for plots of the average graph metric, GCI and CMR_GLU_ in each hub). Older adults had higher GCI for global efficiency in all 13 hubs (except the left and right precentral gyrus and left lateral occipital cortex at the top 30% of edges). Older adults had a higher GCI than younger adults for local efficiency in the contralateral frontal poles and right occipital pole at the top 10% of edges, as well as the middle, superior and inferior frontal gyri at the top 30% of edges. Older adults also had a higher GCI than younger adults for betweenness centrality in the posterior cingulate, precuneus, precentral gyrus and occipital pole at the top 10% of edges. At the top 30% of edges, older adults had a higher GCI for betweenness centrality in the left postcentral gyrus and right inferior frontal gyrus.

**Figure 3.**
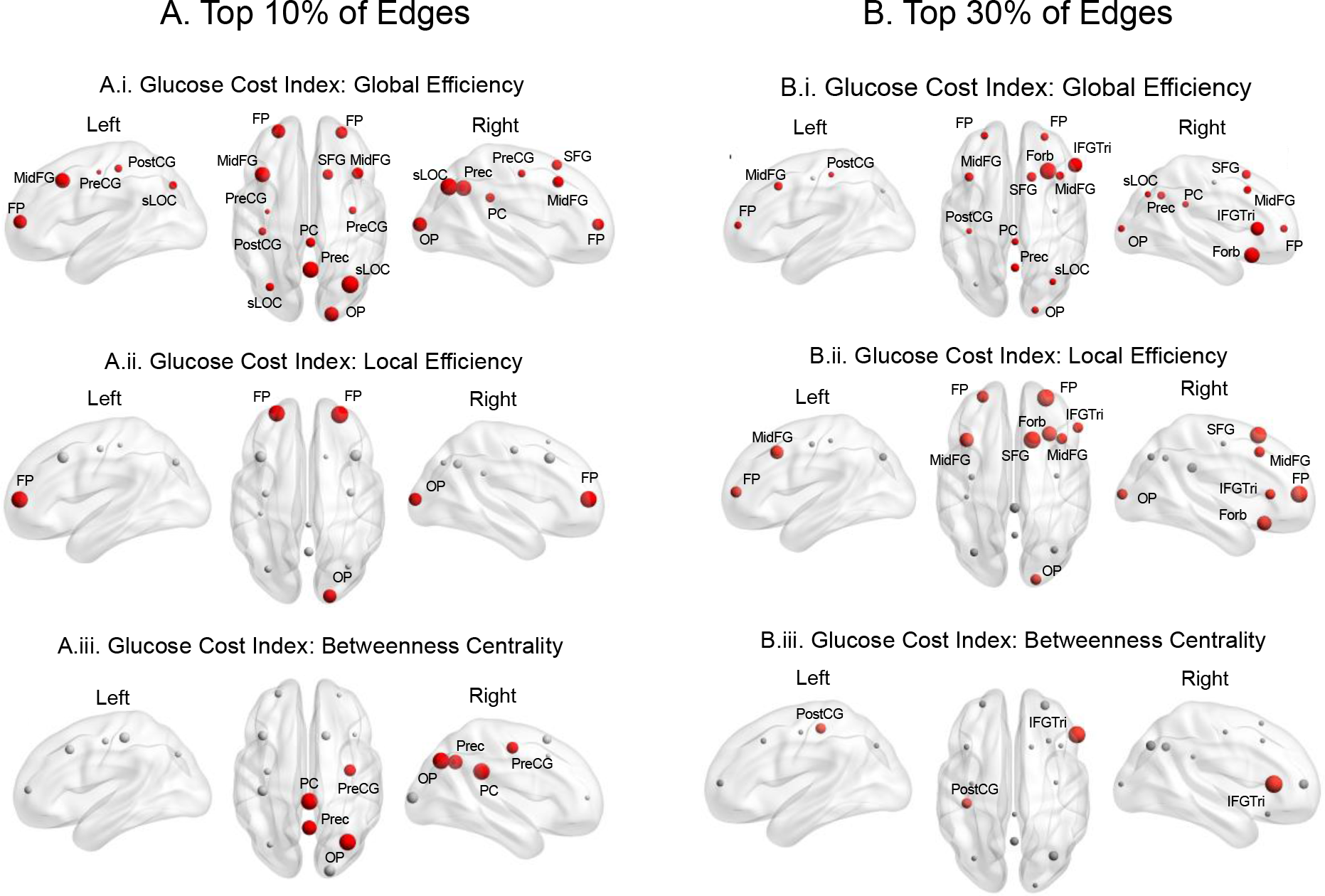
Glucose Cost index (GCI) for hub regions comparing younger and older adults at the top 10% (A) and 30% (B) of network edges. Nodes showing significant age group differences are colored red for older > younger. Nodes showing non-significant age group differences are shaded grey. Relative size of the node reflects the magnitude of the univariate F-statistic following a significant multivariate test (see Supplementary Information, Table S5 and Figure S6). A higher GCI can indicate a more costly use of glucose, stemming from a higher graph measures relative to lower rate or glucose, such as seen in the global efficiency of older adult relative to younger adults. A similar GCI can stem from lower graph measure relative to a lower rate glucose.

### Association Between Metabolic Hub Properties and Cognition

The canonical correlation analysis of global efficiency and cognition identified one significant canonical mode at both the top 10% and 30% of edges (Figure 4). The linear combinations of the global efficiency measures and the cognition measures were significantly correlated with each other, with a correlation coefficient of 0.70 at both the top 10% and 30% of edges (p = 0.013 and p = .048). At the top 10% of edges, higher whole brain and right lateral occipital lobe and precuneus global efficiency (Figure 4A) correlated most strongly with the global efficiency variate and were associated with lower episodic memory, visuospatial processing, cognitive control and response inhibition and slower reaction time (Figure 4B). Conversely, at the top 30% of edges, lower global efficiency for hubs in the frontal and motor regions (Figure 4C) correlated negatively with performance in most cognitive domains (Figure 4D).

**Figure 4.**
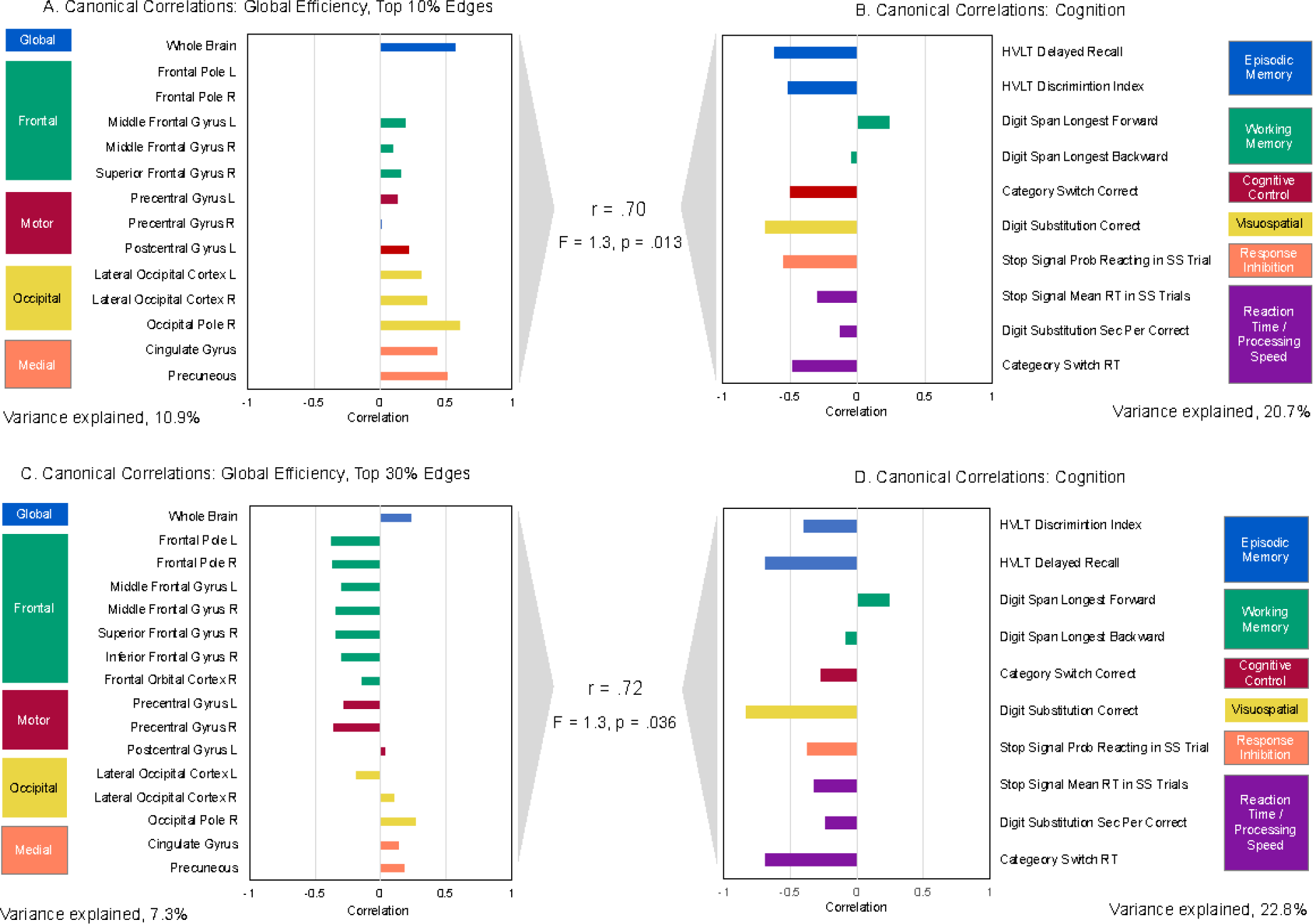
Canonical correlations between the *global efficiency* of the network hubs at (A) top 10% and (B) top 30% of network edges and cognition ((B) and (C)). r-value is the canonical correlation between the linear combinations of the global efficiency and cognition variables that maximally covary across subjects. F-statistic is Wilk’s test of the null hypothesis that the canonical correlation and all smaller ones are equal to zero and was significant for one canonical variate at both the top 10% and 30% of edges. The correlations on each variable set represent the strength of the association between the variable and the canonical variate. Variance explained is the percentage of variance explained by the variables in their variate. Stop signal reaction time, second s per correct response in the digit substitution and category switch reaction time were multiplied by -1 so that higher scores reflect better performance.

The canonical correlation analysis of metabolic hub local efficiency and cognition identified one significant canonical mode at the top 10% of edges and two modes at the top 30% of edges (Figure 5). At the top 10% of edges, lower local efficiency at the whole brain and hub level was associated with worse performance across most cognitive domains (r = .82), most notably episodic memory, visuospatial processing, cognitive control and reaction time and processing speed (Figure 5B). A similar pattern was found at the top 30% of edges for the first canonical factor (r = .77), except that working memory showed the strongest correlation on the cognitive variate. Lower efficiency was associated with lower working memory. The second canonical mode was characterised by lower left and right middle frontal gyri local efficiency, which was associated with lower episodic memory. Interestingly the pattern was opposite for the right superior frontal and cingulate gyri, which correlated positively on the local efficiency variate.

**Figure 5.**
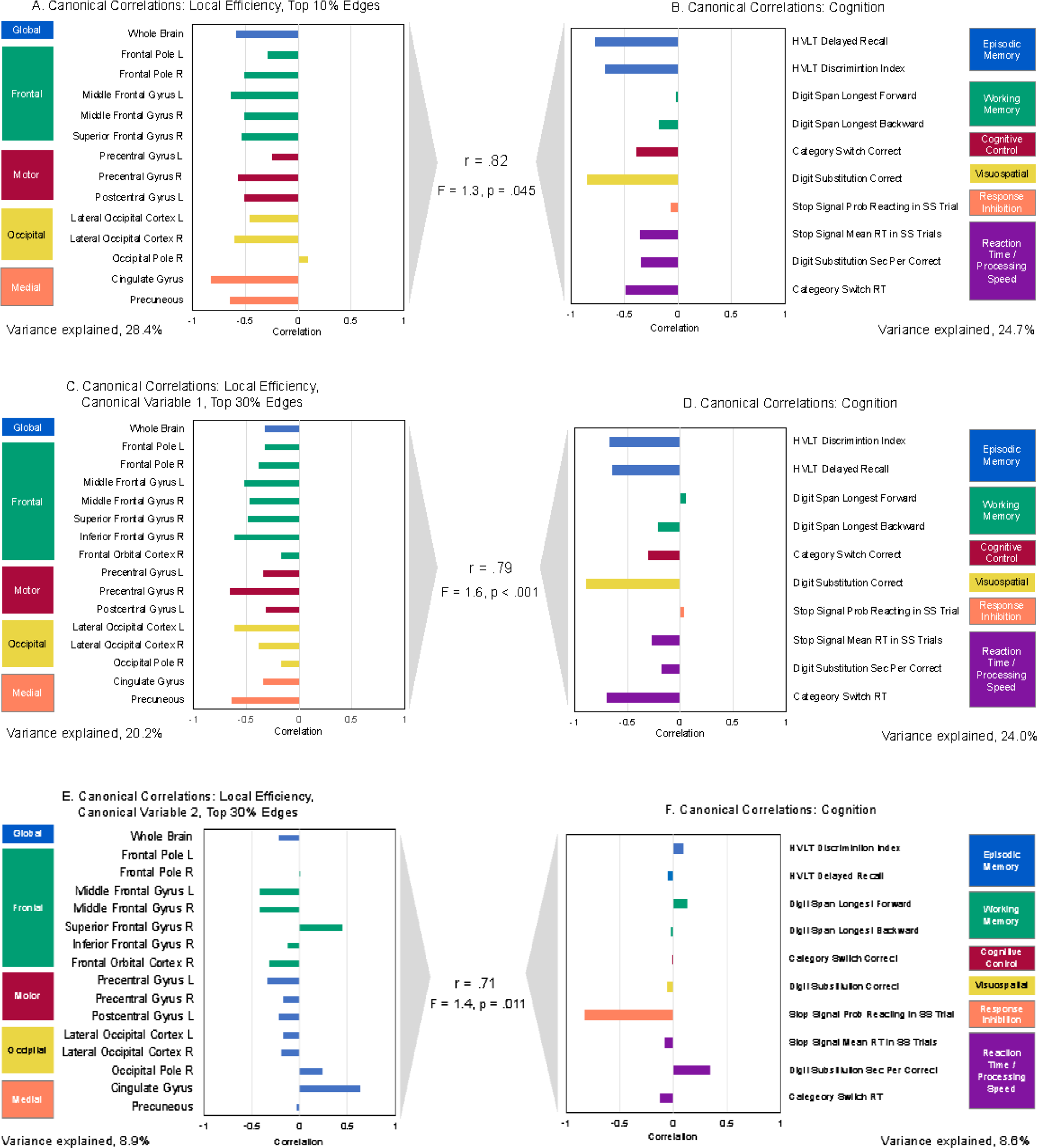
Canonical correlations between the *local efficiency* of the network hubs at (A) top 10% and (B) top 30% of network edges and cognition ((B) and (C)). r-value is the canonical correlation between the linear combinations of the local efficiency and cognition variables that maximally covary across subjects. F-statistic is Wilk’s test of the null hypothesis that the canonical correlation and all smaller ones are equal to zero, and revealed one and two significant canonical variates at the top 10% and 30% of edges, respectively. The correlations on each variable set represent the strength of the association between the variable and the canonical variate. Variance explained is the percentage of variance explained by the variables in their variate. Stop signal reaction time, seconds per correct response in the digit substitution and category switch reaction time were multiplied by -1 so that higher scores reflect better performance.

The canonical correlation analysis of metabolic hub degree and cognition identified one significant canonical mode at the top 10% and 30% of edges (Figure 6). At the top 10% of edges, lower metabolic nodal degree in the frontal regions correlated most strongly on the canonical variate (except the left middle frontal gyrus) and was associated with mostly worse cognitive performance (r = .69). At the top 30% of edges, lower degree in the frontal regions was most associated with worse cognitive control, working and episodic memory and reaction time in the category switch task.

**Figure 6.**
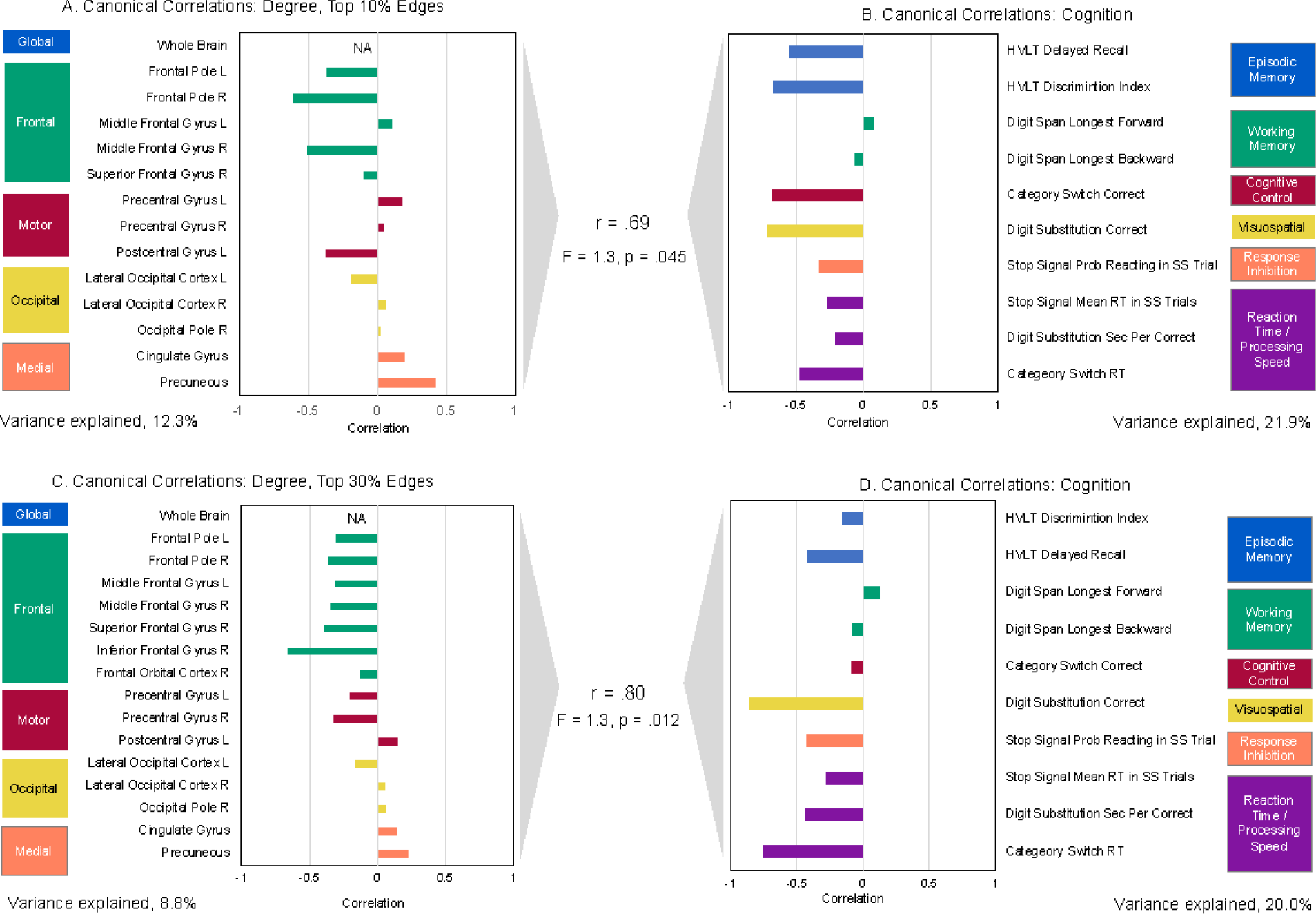
Canonical correlations between *degree* of the network hubs at (A) top 10% and (B) top 30% of network edges and cognition ((B) and (C)). r-value is the canonical correlation between the linear combinations of the degree and cognition variables that maximally covary across subjects. F-statistic is Wilk’s test of the null hypothesis that the canonical correlation and all smaller ones are equal to zero and was significant for one canonical variate at both the top 10% and 30% of edges. The correlations on each variable set represent the strength of the association between the variable and the canonical variate. Variance explained is the percentage of variance explained by the variables in their variate. Stop signal reaction time, seconds per correct response in the digit substitution and category switch reaction time were multiplied by -1 so that higher scores reflect better performance.

No significant canonical correlations were found for betweenness centrality and cognition. None of the canonical correlations were significant controlling for age, i.e., the correlations between the cognition variate and variates of the residuals of the graph metrics after regressing out age differences.

## Discussion

In this first study comparing metabolic connectivity between younger and older adults, we found age related differences in the topology and glucose consumption characterisation of the metabolic connectome. We also found age differences in the efficiency and global integration of metabolic hubs across the brain. Metabolic network hubs had a greater glucose cost than non-hubs, and older people had a lower energy budget to support the connections between hubs and other brain regions. In older people, a sparse metabolic network (top 10% of edges) was more globally integrated, but the integration was driven by stronger connections mainly in the medial and posterior regions; connections that use a larger proportion of a smaller energy budget. However, in a denser network (top 30% of edges), hubs in the frontal regions of older adults showed both a high glucose cost and lower global integration and efficiency than in younger adults, a topology in older adults that was associated with worse cognitive performance. Taken together, these results suggest there are important alterations in the metabolic connectome of older adults, characterised by a reduced capacity of hubs in the frontal regions to connect to other regions, a higher metabolic cost and worse cognitive performance.

Younger and older adults’ metabolic connectomes were distinguished by high connectivity strength for the left and right frontal, motor, parietal and medial cortices, both within and between hemispheres. Consistent with our hypothesis, the largest age differences in metabolic connectivity were in the frontal and temporal regions. These results extend research showing that cerebral metabolic rates of glucose decrease in ageing, particularly in the frontal and temporal regions [7], by indicating that metabolic connectivity in those regions is also the most impacted in ageing.

Since the wiring cost of a region is proportional to nodal degree [24], we studied the metabolic networks at the top 10% and 30% of edges. At these connection densities, we contrasted hub and non-hub regions, to examine if hub regions have a wiring cost premium. Consistent with our hypothesis, we found that, on average, the metabolic hub regions have higher underlying rates of glucose metabolism than non-hub regions and that a higher proportion of the available glucose is used in supporting network connections in hub than non-hub regions. The average glucose cost index was also significantly higher at the top 30% than top 10% of edge for all graph metrics, consistent with the argument that a higher nodal degree and associated connectivity is associated with a higher metabolic cost. These results build on previous research by showing that the glucose cost of metabolic network connectivity increases with increasing connection density. They also complement previous results showing that fMRI network hubs have greater CMR_GLC_ compared to fMRI non-hub regions [40], suggesting that both fMRI and fPET metabolic hubs have a wiring cost premium compared to non-hub regions.

The metabolic network topology of younger and older adults also has implications for cognitive performance. Hubs in the frontal regions were particularly important when canonical correlation analyses were used to test the multivariate relationships between the metabolic hubs and cognition. At the top 30% of edges, higher global and local efficiency in the hubs in the frontal regions, as seen in younger adults, were associated with better cognitive performance and processing speed, including higher episodic memory, visuospatial processing and cognitive control. At the top 10% of edges, higher right lateral occipital global efficiency, as seen in older adults, was associated with lower episodic memory, visuospatial processing, cognitive control, response inhibition and reaction time. These results suggest that the age differences in the global integration of metabolic hubs are at least partly driving cognitive performance. This was further supported when the canonical correlation analyses were repeated controlling for age. The association between graph metrics and cognition were no longer significant, indicating that the age differences in the graph metrics mediated the relationship between metabolic hubs and cognition. Our fPET results are also consistent with the findings from other neuroimaging modalities that hubs are key players in network communication and are prone to age-related alterations that impair cognitive performance [35-37].

Our results support the notion that functional network architecture in the resting state likely reflects at least some of the underlying “circuitry” by which information flows during task performance [41-44]. The functional network properties during a naturalistic viewing setting were associated with cognitive task performance across domains. The fact that frontal, occipital, motor and medial hub regions were related to multiple cognitive domains in the canonical correlation analyses also support the concept of a flexible architecture of the human brain in which functional regions or networks adaptively reconfigure to different behavioural and social context and demands [45, 46].

Our current results extend our previous studies in younger adults [17, 23] by showing that younger and older adults’ metabolic connectomes are distinguished by high metabolic connectivity strength for the frontal, motor, parietal and medial cortices. Here, we used an optimised bolus/infusion radiotracer administration approach [47] and alternative processing pipeline to account for uptake dynamics unrelated to neural activity. Our alternative acquisition and processing pipeline was intended to improve signal to noise, which is lower in the constant infusion approach used in our previous studies [48, 49]. While this approach yielded a more widespread (i.e., beyond frontoparietal regions) and stronger (cf. max r here = 0.4, previously max r = 0.2) pattern of connectivity, the strongest connections were still obtained in frontal, parietal and motor regions.

We have previously reported that older adults have a less modular, more integrated resting-state fMRI functional network architecture, underpinned by lower within-but higher between-network connectivity [31]. Here we found that older adults have a more globally integrated metabolic network but only among the most strongly connected regions in a sparse network (top 10% of egdes). Like the fMRI literature, the metabolic network properties are associated with cognition. However, our results also diverge from the fMRI literature in several ways, including lower metabolic connectivity in older adults between the frontal and temporal regions and other lobular regions, and the most efficient regions in older adults being mostly in the medial and posterior regions of the brain. This contrasts with the fMRI literature which shows increased connectivity strength between networks nodes in the frontal and temporal regions and a posterior to anterior shift in ageing (see [31] for review).

The reason for these differences remains to be determined. If we accept that temporal coherence of regional functional signals reflects information transfer within the brain [16, 50], then our results suggest that there is reconfiguration of the metabolic signals in ageing. While both the metabolic and hemodynamic signals reflect functional brain activity, they do in fact measure different aspects of neuronal activity. Increased glucose metabolism contributes to the haemodynamic response measured by BOLD-fMRI, however the BOLD response also requires a regional increase in cerebral blood flow, metabolic rate of oxygen, lactate, and a decrease in oxygen extraction fraction [51] to be detected. Ageing affects each of these indices independently and variably across the brain [7, 52-54]. By contrast, FDG-PET is considered a less confounded measure of glucose uptake at the excitatory post-synaptic neuron [55, 56]. As such, differences in metabolic and functional fMRI connectivity are not surprising, and likely reflect different effects of ageing on the cascade of events that contribute to the BOLD response. Taken together, our results are compatible with our previous conclusion that metabolic connectivity indexes unique and complementary aspects of information transfer in the brain compared to fMRI connectivity [17, 23].

One limitation of this study is the cross-sectional design, which limits conclusions about the causal relationships between age, metabolic network properties and cognition. Longitudinal research can directly test whether changes in metabolic networks in younger and older individuals lead to changes in cognition. Our study is also limited by the absence of middle age adults. Research on structural and functional brain networks using fMRI have reported quadratic trajectories of age differences, with an inflection point somewhere in the third to fifth decade of life (see [31] for review). However, the lack of middle aged adults precludes the identification and quantification of ageing trajectories of the metabolic networks across the full adult lifespan. Additional research is needed to study these trajectories.

As demonstrated here, advances in fPET methods have led to improvements in the temporal resolution and signal-to-noise. Nonetheless, the methods are still underdeveloped relative to the well-established standard PET and fMRI processing pipelines and analysis methods. PET reconstruction and filtering methods [49], including machine learning algorithms [57], may further increase sensitivity of functional measurement. Methods to quantify the resting-state fPET signal are still under development. Importantly, fPET is not the only approach that can be used to measure within-subjects PET connectivity [39] and it remains to be determined how complementary these approaches are in estimating metabolic connectivity. In addition, with improved methods, it may be possible to reduce the frame duration to a temporal resolution approaching fMRI and to index metabolic network properties at faster temporal frequencies. Such research may shed new light on the metabolic networks at rest and during task performance.

In conclusion, using high temporal resolution fPET we show for the first time that there are age differences in the metabolic connectivity and glucose cost of global network integration in younger and older adults. Our results build on work in other imaging modalities suggesting that metabolic hubs are of particular importance for optimal brain function because they support information integration across the brain. Ageing is characterised by an altered metabolic network topology and reduced capacity of hubs in the frontal regions to connect to other regions. The hubs of younger and older adults use a high proportion of the available glucose in the brain to support global integration relative to non-hub regions, presumably reflecting a high cost of glucose metabolism in the synaptic activity of the hubs. Younger adults use a high proportion of their glucose metabolism budget to support the efficiency and global integration of hubs in the frontal regions. In contrast, older adults use a high proportion of their smaller energy budget to support global integration of mostly posterior and medial hub regions, a pattern associated with worse cognitive performance. These results highlight the fundamental role that metabolism plays in supporting information transfer in the brain, especially in hub regions, and the unique insights that metabolic connectivity provides into the ageing brain.

## Method

Additional details of the methods are provided in the Supplement.

### Ethical Considerations

The study was approved by the Monash University Human Research Ethics Committee. Participants provided informed consent to participate in the study.

### Participants

Ninety participants were recruited from the local community. An initial screening interview ensured that participants did not have a history of diabetes, neurological or psychiatric illness and were not taking psychoactive medication. Participants were also screened for claustrophobia, non-MR compatible implants and a clinical or research PET scan in the previous 12 months. Women were screened for current or suspected pregnancy. Participants received a $100 voucher for participating in the study. Four participants were excluded for further analyses due to either excessive head motion (N=2) or incomplete PET (N=2) data.

### Data Acquisition

Participants completed an online demographic and lifestyle questionnaire and a cognitive test battery. Briefly, the following cognitive measure were used (see Supplement for details): delayed recall and a recognition discrimination index from the Hopkins Verbal Learning Test; length of longest correct series of forward and backward recall from a digit span test to index working memory; the proportion of correct trials and mean reaction time in a task switching test to index cognitive control and flexibility; the probability of responding and reaction time in a stop signal task to measure response inhibition; and number of correct responses and seconds per correct response in a digit substitution task to measure visuospatial performance and processing speed.

Participants underwent a 90-minute simultaneous MR-PET scan in a Siemens (Erlangen) Biograph 3-Tesla molecular MR scanner. At the beginning of the scan, half of the 260 MBq FDG tracer was administered via the left forearm as a bolus, the remaining 130 MBq of the FDG tracer dose was infused at a rate of 36ml/hour over 50 minutes [47]. Non-functional MRI scans were acquired during the first 12 minutes, including a T1 3DMPRAGE and T2 FLAIR. Thirteen minutes into the scan, list-mode PET and T2* EPI BOLD-EPI sequences were initiated. A 40-minute resting-state scan was undertaken in naturalistic viewing conditions. Full details of the acquisition parameters are provided in Supplementary Information Section 1. Here, we focus on results from the PET acquisition.

### MRI and PET Pre-Processing

For the T1 images, the brain was extracted in Freesurfer; quality of the pial/white matter surface was manually checked, corrected and registered to MNI152 space using Advanced Normalization Tools (ANTs). The list-mode PET data for each subject were binned into 344 3D sinogram frames of 16s intervals. Attenuation was corrected via the pseudo-CT method for hybrid PET-MR scanners [58]. Ordinary Poisson-Ordered Subset Expectation Maximization algorithm (3 iterations, 21 subsets) with point spread function correction was used to reconstruct 3D volumes from the sinogram frames. The reconstructed DICOM slices were converted to NIFTI format with size 344 × 344 × 127 (size: 1.39 × 1.39 × 2.03 mm^3^) for each volume. All 3D volumes were temporally concatenated to form a single 4D NIFTI volume. After concatenation, the PET volumes were motion corrected using FSL MCFLIRT [59], with the mean PET image used to mask the 4D data. PET images were corrected for partial volume effects using the modified Müller-Gartner method [60] implemented in Petsurf.

### Metabolic Connectomes

Participants’ T1 images and pre-processed fPET timeseries data were loaded to the CONN toolbox in Matlab [61]. The first 10 minutes of the timeseries was excluded to remove the initial radiotracer uptake and to reflect the time corresponding to the onset of a stable signal [48]. The timeseries was denoised using the default CONN pipeline, regressing out the potential confounding from white matter and CSF. A low pass filter (.0625 Hz) was applied to filter out high frequency noise in the PET signal [62]. Regions of interest were generated for 106 cortical and subcortical regions of the Harvard Oxford atlas, excluding the cerebellum. As it is currently unclear if fMRI-derived ‘functional’ atlases, such as the Schaefer atlas [63], are transferable to metabolic connectomes [17, 19, 20, 39], we applied an anatomical parcellation to this data. However, see Supplementary Information Section 2.1 for discussion of atlas selection and results for the Schaefer 100 functional atlas.

The following graph theory metrics were calculated in CONN: Global efficiency, local efficiency, betweenness centrality and degree (see Supplement for definitions). *Metabolic hubs* were defined as regions more than one standard deviation above the mean on at least two of betweenness centrality, degree and global efficiency. We adopted this definition because no single metric unambiguously defines a node as a hub [33, 37].

### CMR_GLU_ and Glucose Cost Index

Calculations of regional CMR_GLU_ were undertaken in Magia [64] using the FDG time activity curves for the Harvard Oxford atlas regions. The FDG in the plasma samples was decay-corrected for the time between sampling and counting as the input function to Patlak models. The FDG and plasma data were used from the 10 minute point onwards. Each participant’s baseline plasma glucose (mmol) was entered in the Patlak model. Regional CMR_GLU_ could not be calculated for three younger and two older participants due to blood haemolysis or well counter issues. Those participants were excluded from analyses in which CMR_GLU_ was used.

We computed the Glucose Cost Index (GCI) as the ratio between each graph metric and CMR_GLU_ in each hub region. The GCI values reflect the metabolic cost of global efficiency, local efficiency, betweenness centrality and degree.

### Analyses of Age Group Differences

#### Metabolic Connectomes

To test the first hypothesis of widespread connectivity beyond the frontoparietal regions, we qualitatively assessed the metabolic connectomes of older and younger adults. The metabolic connectomes were generated by reverse transforming the individual participant matrices of Fisher transformed z-values back to r-values, and calculating the average for each node separately for the younger and older adult groups.

#### Within and Between Region Connectivity

To quantify the connectivity in the younger and older adult connectomes, within- and between region connectivity was calculated. *Within connectivity* was calculated as the average Fisher transformed z-value of all regions within the frontal, temporal, medial temporal, parietal, occipital, motor, limbic, medial and subcortical lobes / major cortical areas. *Between connectivity* was calculated as the average of the regional connections within that lobe or major cortical area to the regions in all other lobes or major cortical areas. We used independent sample *t*-tests (p-FDR < .05, one tailed) to test the hypothesis that older adults will have lower within- and higher between connectivity than younger adults, particularly in the frontal and temporal regions.

### Function of Metabolic Hub and Non-Hub Regions

A series of multivariate general linear models (GLMs) was used to test the third hypothesis that older adults will show higher network integration and lower efficiency and hub function . To control Type I error rates, the graph metrics for all of the metabolic hub regions were used as the dependent variables in separate multivariate GLMs with age group as the predictor. Each GLM was tested at p<.05 uncorrected. Where an overall multivariate GLM was statistically significant, separate follow-up univariate tests of age group differences for each hub were undertaken.

Separate analyses were undertaken at the top 10% and 30% of network edges. We chose the 10% and 30% edge thresholds for two reasons. First, graph metrics, particularly global efficiency, are sensitive to network degree as it alters the number and length of edges between nodes [65]. Second, the “wiring cost” of functional fMRI connectivity increases monotonically with increasing connection density [24]. In fMRI networks, between approximately 5% and 34% of edges in a network of 90 nodes maximises the efficiency of the network topology in proportion to their connection cost [29]. By using two edge thresholds, we examined the trade-off in the metabolic network efficiency and cost between the edge thresholds, with an assumed higher wiring cost at the top 30% of edges. We also assessed whether hub function and topology are differentially associated with cognitive performance at the different thresholds. Measures from canonical correlation analyses were used to quantify these relationships (described below).

To further characterise the metabolic networks, younger and older adults were also compared on whole brain global efficiency, local efficiency, betweenness centrality and degree (i.e., for the entire network) and at each of the 106 nodes using independent sample *t*-tests (p-FDR < .05, two-tailed, for each series of analyses).

### Cerebral Metabolic Rates of Glucose and Glucose Efficiency Index of Hub and Non-Hub Regions

For each participant, the percentage CMR_GLU_ in the hub regions relative to the non-hub regions was calculated by dividing the average rate in the hubs by the average rate in the non-hubs. A one-sample *t*-test was used to test whether the percentage difference was significantly greater than zero.

To test the hypothesis that a greater proportion of glucose metabolism will be used to support network connections in the hub than non-hub regions, the average GCI of the hubs and non-hub regions at the 10% and 30% edge thresholds were compared in separate repeated GLMs across the entire sample for each graph metric. Finally, multivariate GLMs were used to test for differences between older and younger adults in the GCI for each graph metric in the hub regions. Following the presence of a statically significant multivariate model, separate univariate tests comparing older and younger adults were undertaken for each hub region.

### Association Between Metabolic Hub Properties and Cognition

To test the hypothesis that higher efficiency, degree and centrality of metabolic hubs will be associated with better cognitive performance, we ran a series of canonical correlational analyses using the 10 cognitive measures and the graph theory metrics for the hub regions. Stop signal reaction time, seconds per correct response in the digit substitution and category switch reaction time were first multiplied by -1 so that higher scores reflect better performance. The cognitive scores were then converted to Z-scores. The graph theory measure at the whole brain and in the metabolic hubs at the top 10% and 30% of edges were included in separate canonical correlation analysis to return a cognitive profile associated with the graph measures. The number of potential canonical variates in each analysis reflects the smallest number of variables in either set, in this case the 10 cognitive measures. The F-statistic from the Wilk’s test was used to test the null hypothesis that the canonical correlation and all smaller ones are equal to zero. To aid interpretation of the variables most contributing to the canonical variates, we focus on correlation coefficients ≤ −0.21 and ≥0.21 between the variable and the variate.

To assess whether age mediates the relationship between cognition and the metabolic networks, the canonical correlation analyses controlling for age group were repeated. First, age group was regressed onto each graph metrics at each hub region and the residuals saved for each participant. Canonical correlation analysis was then undertaken between the cognition measures and the residuals of the graph metrics.

## Data Availability Statement

The datasets used and/or analysed during the current study available from the corresponding author on reasonable request.

## Acknowledgements

Jamadar is supported by an Australian National Health and Medical Research Council (NHMRC) Fellowship (APP1174164).

## Author Contributions

SJ conceptualized and designed the study. HD, MNS, GM, RDP collected the data, and with KV & EXL prepared the data for analysis. HD and SJ designed the analyses and manuscript. HD and EXL undertook the analyses. HD wrote the first draft of the manuscript with SJ edits. CM & GFE provided supervision. All authors contributed to manuscript preparation or review. We thank Richard McIntyre, Lauren Hudswell and the staff at Monash Biomedical Imaging for their contributions to data acquisition and image reconstruction.

## Footnotes

1 Here, we reserve the term ‘metabolic connectivity’ to refer to a within-subjects measurement of the coherence of time-varying regional FDG signals (reflecting glucose uptake), and the term ‘metabolic covariance’ to refer to across-subject correlation of regional FDG signals. See ref (17) for discussion.

