## Supplementary Information for "Metabolic connectivity in ageing"

~

### 1. Supplementary Methods

#### 1.1 Study Ethics

The study protocol was reviewed and approved by the Monash University Human Research Ethics Committee, in accordance with Australian Code for the Responsible Conduct of Research (2007) and the Australian National Statement on Ethical Conduct in Human Research (2007). Participants provided informed consent to participate in the study. Administration of ionizing radiation was approved by the Monash Health Principal Medical Physicist, following the Australian Radiation Protection and Nuclear Safety Agency Code of Practice (2005). For participants older than 18 years, the annual radiation exposure limit of 5 mSv applies; the effective dose in this study was 4.9 mSv.

#### 1.2 Cognitive Battery

Prior to the scan, participants completed an online demographic and lifestyle questionnaire including age, sex, education, height and weight, history of smoking, alcohol and recreational drug use. Participants also completed a cognitive test battery consisting of measures of general intelligence, working memory, cognitive flexibility, inhibitory control and verbal learning.

**Wechsler Abbreviated Scale of Intelligence (WASI-IQ).** An assessment of intelligence suitable for ages 6-90 years [1]. There are 4 subtests: block design, vocabulary, matrix reasoning and similarities. WASI-IQ was scored by converting raw scores into a scale score, which were transformed into a composite score reflecting verbal comprehension and perceptual reasoning abilities (FSIQ2). This score was converted to an age-based T scores established in a normal population.

**Hopkins Verbal Learning Test (HVLT).** A three-trial list learning and free recall task comprising 12 words, four words from each of three semantic categories [2]. Approximately 20–25 minutes later, a delayed recall trial and a recognition trial was completed. The delayed recall required free recall of any words remembered. The recognition trial comprised 24 words, including the 12 target words and 12 false-positives, six semantically related, and six semantically unrelated. Delayed recall (total words recalled) and a recognition discrimination index (number of correct minus number of false positives in the recognition task) were calculated.

**Digit Span.** A measure of verbal short term and working memory used in two formats: Forward and backward digit span [3]. Participants were presented with a series of digits, and are asked to repeat them in either the order presented (forward span) or in reverse order (backwards span). After two consecutive failures of the same length, the test was stopped. Scores were derived as the length of longest correct series for both forward and backward recall.

**Task Switching.** A computer-based test in which participants were given a word and had to perform one of two simple categorisation tasks, depending on the cue that appeared with the word: 1) 'living' task. If the cue was a heart, participants were asked to categorise the word via a key press based on whether it represents a LIVING versus a NON-LIVING object; and 2) 'size' task. If the cue was an arrow-cross, participants were asked to categorise the word via a key press based on whether it represents an object that is BIGGER or SMALLER than a basketball. The cue selection for each new trial was randomised. Half the test trials were switch trials; half non-switch trials. Half the switch and non-switch trials were congruent in the key presses for either task, half were incongruent. The measures used included the mean latency of correctly responding to a switch trial and switch cost. Switch cost is the difference between mean correct latency of switch trials and nonswitch trials with positive value indicating participants were slower on switch trials, that is, there was a latency cost to switching [4].

**Stop Signal.** A computer-based test in which participants were presented an arrow that pointed either right or left [5]. The task was to press the left response key if the arrow pointed to the left and press the right response key if the arrow pointed to the right, unless a signal beep is played after the presentation of the arrow. In this case the response should be stopped before execution. The delay between presentation of arrow and signal beep (starting at 250ms) was adjusted up or down (by 50ms) depending on performance. The delay got longer if the previous signal stop was successful (up to 1150ms) and smaller if the previous signal stop was not successful (down to 50ms). The stimulus onset asynchrony between the start of each trial (onset of fixation circles) was 2000ms. Variables were the mean reaction time in stop signal trials and stop signal reaction time. Stop signal reaction time is an estimate of inhibition ability, that is, the time required to stop the initiated go-process. The slower the stop signal reaction time, the more difficult to stop the go-process.

**Digit Symbol Substitution.** A computer-based task in which participant were presented with an 18 columns x 16 rows matrix [6]. The task was to translate symbols shown above the matrix (key) into digits in the matrix within a two minute period. Total count of correct responses and seconds per correct response were recorded.

#### 1.3 MR-PET Data Acquisition

Participants underwent a 90-minute simultaneous MR-PET scan in a Siemens (Erlangen) Biograph 3-Tesla molecular MR scanner. Participants were directed to consume a high-protein/low-sugar diet for the 24 hours prior to the scan. They were also instructed to fast for six hours and to drink 2–6 glasses of water. Prior to FDG infusion, participants were cannulated in the vein in each forearm and a 10ml baseline blood sample taken. At the beginning of the scan, half of the 260 MBq FDG tracer was administered via the left forearm as a bolus, providing a strong PET signal from the beginning of the scan. The remaining 130 MBq of the FDG tracer dose was infused at a rate of 36ml/hour over 50 minutes, minimising the amount of signal decay over the course of the data acquisition. We have previously demonstrated that this protocol provides a good balance between a fast increase in signal-to-noise ratio at the start of the scan, and maintenance of signal-to-noise ratio over the duration of the scan [7].

Participants were positioned supine in the scanner bore with their head in a 32-channel radiofrequency head coil and were instructed to lie as still as possible. The scan sequence was as follows. Non-functional MRI scans were acquired during the first 12 minutes, including a T1 3DMPRAGE (TA = 3.49 min, TR = 1640ms, TE = 234ms, flip angle = 8°, field of view = 256 × 256 mm<sup>2</sup>, voxel size = 1.0 × 1.0 × 1.0 mm<sup>3</sup>, 176 slices, sagittal acquisition) and T2 FLAIR (TA = 5.52 min, TR = 5,000ms, TE = 396ms, field of view = 250 × 250 mm<sup>2</sup>, voxel size = .5 × .5 × 1 mm<sup>3</sup>, 160 slices) to image the anatomical grey and white matter structures, respectively. Thirteen minutes into the scan, list-mode PET (voxel size = 1.39 × 1.39 × 5.0mm<sup>3</sup>) and T2\* EPI BOLD-fMRI (TA = 40 minutes; TR = 1000ms, TE = 39ms, FOV = 210 mm<sup>2</sup>, 2.4 × 2.4 × 2.4 mm<sup>3</sup> voxels, 64 slices, ascending axial acquisition) sequences were initiated. A 40-minute resting-state scan was undertaken in naturalistic viewing conditions watching a movie of a drone flying over the Hawaii Islands. At 53 minutes, pseudo-continuous arterial spin labelling (pc-ASL) began, and at 58 minutes, diffusion-weighted imaging (DWI) was acquired with 71 directions to index white matter connectivity. pcASL, DWI and fMRI results are not reported here.

Plasma radioactivity levels were measured throughout the duration of the scan. Beginning at 10-minutes post infusion onset, 5ml blood samples were taken from the right forearm using a vacutainer at 10-minute intervals for a total of nine samples. The blood sample were immediately placed in a Heraeus Megafuge 16 centrifuge (ThermoFisher Scientific, Osterode, Germany) and spun at 2,000 rpm (RCF ~ 515g) for 5 minutes. 1,000-μL plasma was pipetted, transferred to a counting tube, and placed in a well counter for four minutes. The count start time, total number of counts, and counts per minute were recorded for each sample.

##### 1.4 Correction for Partial Volume Effects

PET images were corrected for partial volume effects using the modified Müller-Gartner method implemented in PetSurf (https://surfer.nmr.mgh.harvard.edu/fswiki/PetSurf) [8, 9]. The method corrects for white matter spill in and grey matter spill out of the PET signal. The equation subtracts from the grey matter voxel signal the white matter signal (convoluted by the point spread function) and divides by the grey matter signal. This division can introduce over-correction at the grey matter boundary, and hence a grey matter binary mask is recommended, with the threshold level needing to be chosen. A grey matter threshold of 20-30% is recommended in ageing because atrophy can influence results [8]. For our analyses, we chose a 25% grey matter threshold and surface-based spatial smoothing [9]. We used a Gaussian kernel with a full width at half maximum of 8mm to increase the signal-to-noise ratio. Subcortical structures were partial volume corrected and spatially smoothed in volume space and merged with the cortical data.

##### 1.5 Graph Theory Metrics

The following graph theory metrics were calculated:

**Global Efficiency** at each node defined as the average of the shortest inverse-distances between the node and all other nodes in the graph. Across the entire graph, global efficiency represents a measure of global integration.

**Local Efficiency** at each node defined as the average of shortest inverse-distances between the nodes within the neighbouring sub-graph (all nodes neighbouring that node and all existing edges among them). Network local efficiency is a measure of local integration of a network.

**Betweenness Centrality** defined as the proportion of times that a node is part of a shortest-path between any two nodes within a graph. It represents a measure of node centrality within a graph.

**Degree** at each node defined as the number of edges from and to that node. Degree characterises the local connectedness of each region within the network.

### 2. Supplementary Results

#### 2.1 'Functional' Parcellation

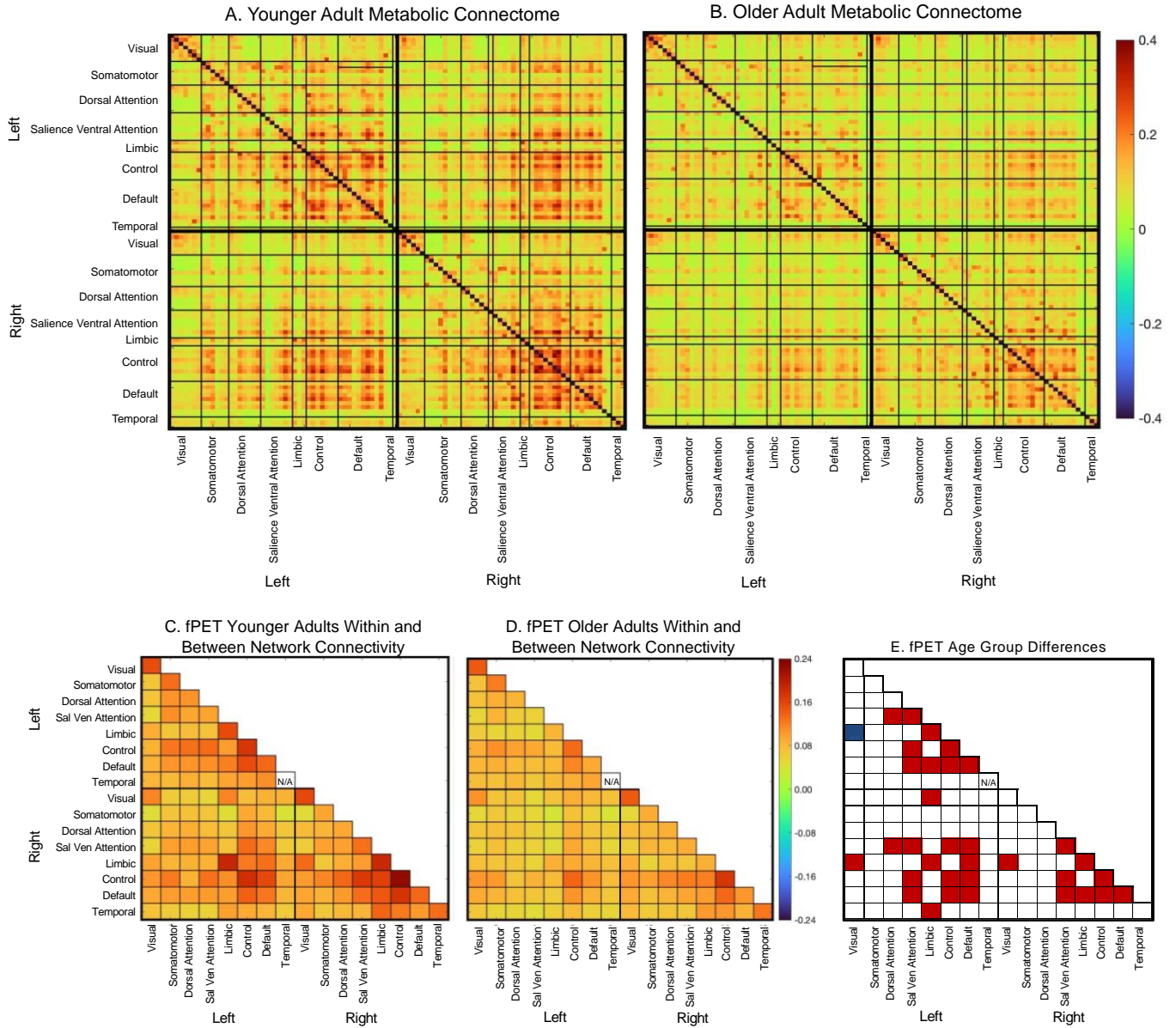

**Figure S1.** Metabolic connectivity for younger and older adults using the Schaefer parcellation. (A) Younger and (B) older adult metabolic connectomes across the 100 nodes in 8 functional networks. Within and between network connectivity averaging the nodes within the networks for (C) younger and (D) older adults. (E) significance test (T-test, $df = 84$) of younger vs older adults, with red shading indicating younger > older and blue shading older > younger differences at $p\text{-FDR} < .05$. Note: N/A for within left temporal network reflects one network region in the Schaefer parcellation.

There is currently no consensus on the optimal atlas to parcellate metabolic data. In fMRI, it is widely accepted that a 'functional' atlas derived from resting-state fMRI data provides a robust estimates and high interpretability of functional connectivity results (e.g., Schaefer atlas [10]). Brain parcellations derived from a single modality (e.g., resting-state fMRI) show varying levels of transferability to other modalities [11] and it is currently untested how transferable fMRI-derived atlases are to FDG-PET data. Previous results from metabolic covariance analyses [12, 13] indicate that fMRI-derived atlases are probably not directly transferable to FDG-PET data. Therefore, we chose to use an anatomical parcellation (and sort) for results in the main paper. However, results for the Schaefer 100 atlas are provided here for qualitative comparison. In addition, because PET SNR scales directly with spatial scale and the size of the ROI [14], we chose parcellations with relatively coarse granularity.

The most salient characteristic for the group-averaged metabolic connectome based on the Schaefer parcellation for both younger and older adults was high connectivity strength in and between the control and default networks, both within and between hemispheres (Fig. S1A and S1B). Sensorimotor, visual, dorsal and salience ventral attention with network connectivity were also evident for the younger and older adults. In contrast, connectivity strength was relatively low in the limbic and left temporal networks. A second salient feature is the apparent lower connectivity strength across the connectome for older than younger adults, as seen by less dark red-to-orange.

### 2.2 Definition of Metabolic Hubs

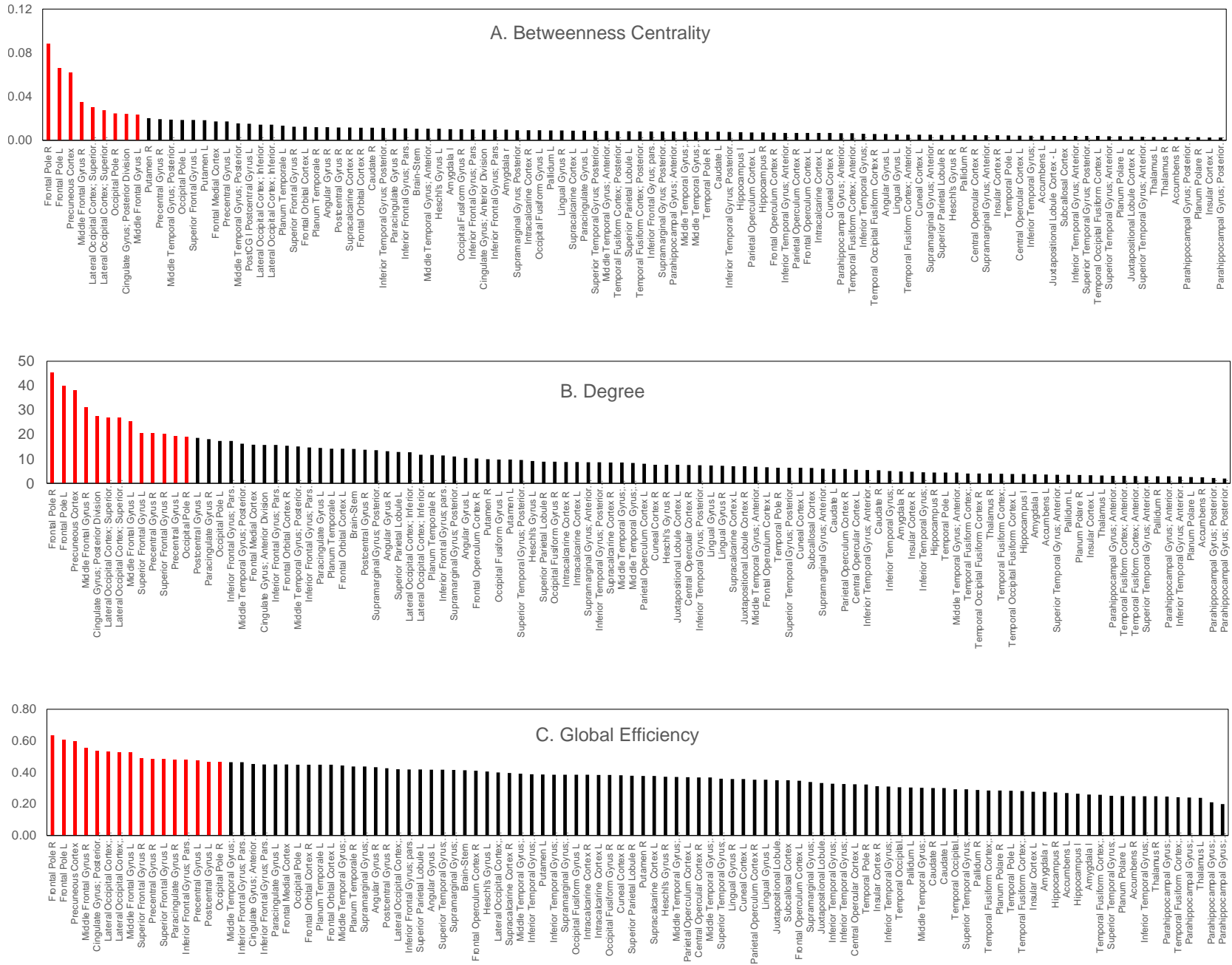

**Figure S2.** Rank order of average (A) Betweenness Centrality, (B) Degree, and (C) Global Efficiency across all subjects at top 10% of edges. Red bars indicate nodes > 1 standard deviation above the mean. Nodes > 1 SD on at least two measures were defined as metabolic hubs. The regions included the left and right frontal poles, bilateral middle frontal gyri and right superior frontal gyrus in the frontal lobes; the left and right precentral gyri and left postcentral gyri in the motor cortex; the bilateral superior division of the lateral occipital cortices and the right occipital pole in the occipital lobe; and the posterior division of the cingulate gyrus and precuneus in the medial cortex.

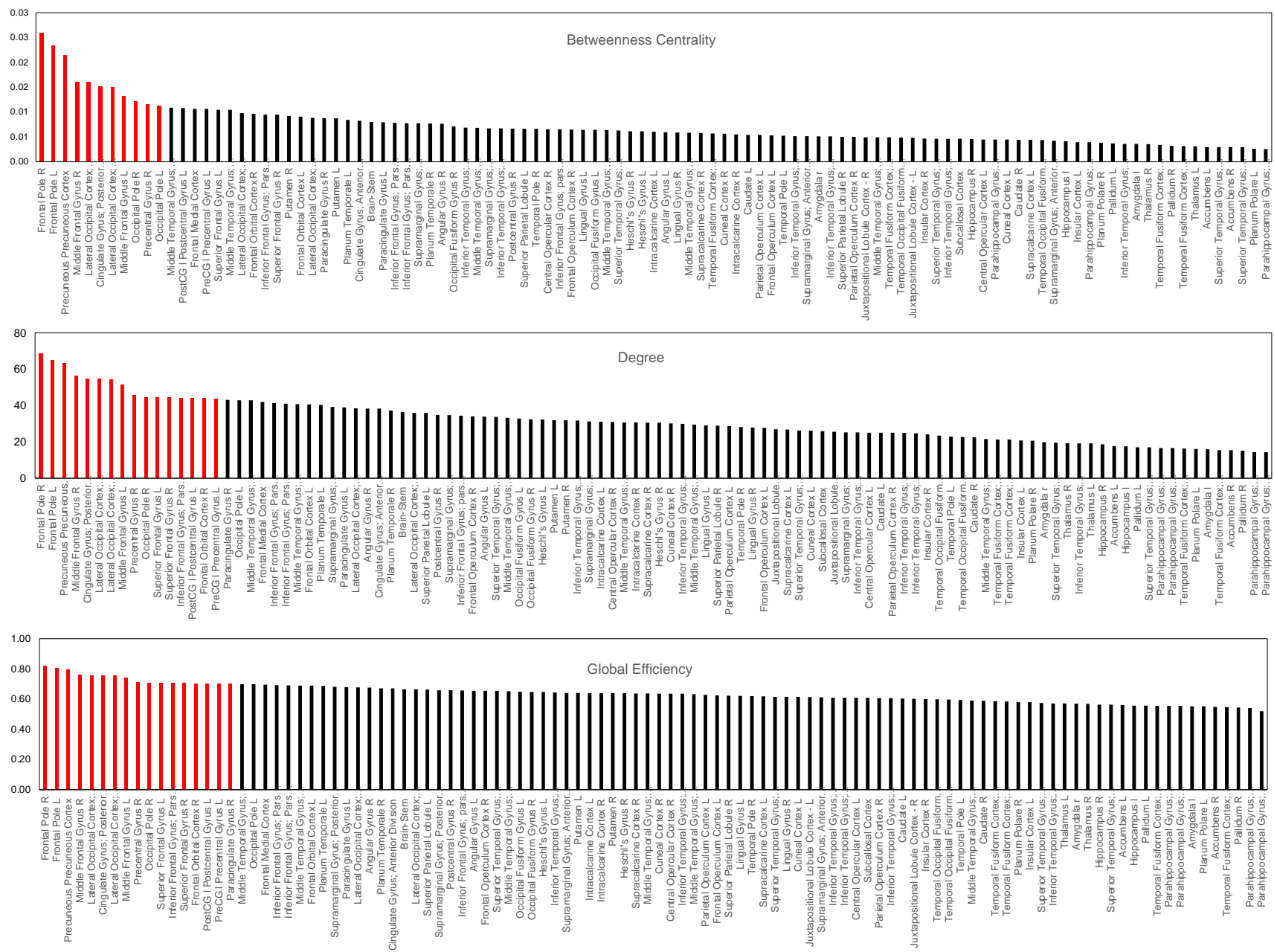

**Figure S3.** Rank order of average (A) Betweenness Centrality, (B) Degree, and (C) Global Efficiency across all subjects at top 30% of edges. Red bars indicate nodes > 1 standard deviation above the mean. All nodes > 1 SD on at least two measures were also identified as hubs at top 10% of edge (see Figure S2), with the addition of the right Inferior Frontal Gyrus, Pars Triangularis; and the right Frontal Orbital Cortex.

### 2.3 Age Differences in the Function of Metabolic Hubs

The mean and standard deviation for older and younger adults on the graph metrics in the metabolic hubs are shown in Table S1. Age group differences were found in the multivariate test at the top 10% of edges for global efficiency ($F(15,70) = 2.0$, $p < .05$), local efficiency ($F(15,70) = 3.4$, $p = .001$), betweenness centrality ($F(15,70) = 2.9$, $p = .002$) and degree ($F(15,70) = 2.7$, $p = .003$). Age differences were also found at the top 30% of edges for global efficiency ($F(15,70) = 13.6$, $p < .001$), local efficiency ($F(15,70) = 4.4$, $p < .001$), betweenness centrality, ($F(15,70) = 3.5$, $p < .001$) and degree ($F(15,70) = 3.0$, $p = .001$). The follow-up univariate F-test of age group differences in the metabolic hubs are shown in Table S1

**Table S1.** Mean, Standard Deviation (SD) and post-hoc univariate F-test (df = 1,84) of age group differences in graph metrics<sup>1</sup> for metabolic hubs at top 10% and 30% of network edges. F-values are plotted on the glass brains in Figures 2 in the main manuscript.

|  | Global Efficiency |  |  |  |  |  | Local Efficiency |  |  |  |  |  | Betweenness Centrality |  |  |  |  |  | Degree |  |  |  |  |  |
| --- | --- | --- | --- | --- | --- | --- | --- | --- | --- | --- | --- | --- | --- | --- | --- | --- | --- | --- | --- | --- | --- | --- | --- | --- |
|  | Younger |  | Older |  | Younger Vs Older |  | Younger |  | Older |  | Younger Vs Older |  | Younger |  | Older |  | Younger Vs Older |  | Younger |  | Older |  | Younger Vs Older |  |
|  | Mean | SD | Mean | SD | F | p | Mean | SD | Mean | SD | F | p | Mean | SD | Mean | SD | F | p | Mean | SD | Mean | SD | F | p |
| Top 10% of Edges |  |  |  |  |  |  |  |  |  |  |  |  |  |  |  |  |  |  |  |  |  |  |  |  |
| Frontal Pole L | 0.60 | 0.06 | 0.61 | 0.07 | 0.5 | 0.000 | 0.69 | 0.07 | 0.65 | 0.10 | 2.9 | 0.093 | 0.068 | 0.037 | 0.063 | 0.033 | 0.4 | 0.512 | 43.1 | 8.9 | 37.1 | 10.6 | 8.1 | 0.006 |
| Frontal Pole R | 0.63 | 0.07 | 0.64 | 0.06 | 0.2 | 0.000 | 0.66 | 0.06 | 0.62 | 0.06 | 5.1 | 0.028 | 0.098 | 0.049 | 0.080 | 0.039 | 3.7 | 0.058 | 48.9 | 8.4 | 42.3 | 10.2 | 10.5 | 0.002 |
| Middle Frontal Gyrus L | 0.53 | 0.06 | 0.52 | 0.07 | 0.0 | 0.000 | 0.76 | 0.05 | 0.67 | 0.15 | 11.5 | 0.001 | 0.022 | 0.025 | 0.025 | 0.024 | 0.4 | 0.523 | 29.1 | 10.3 | 21.8 | 10.1 | 10.9 | 0.001 |
| Middle Frontal Gyrus R | 0.55 | 0.11 | 0.55 | 0.08 | 0.0 | 0.000 | 0.73 | 0.07 | 0.64 | 0.15 | 8.6 | 0.005 | 0.039 | 0.032 | 0.031 | 0.023 | 1.6 | 0.205 | 35.0 | 9.0 | 27.5 | 12.1 | 10.2 | 0.002 |
| Superior frontal Gyrus R | 0.49 | 0.09 | 0.48 | 0.08 | 0.1 | 0.000 | 0.81 | 0.08 | 0.63 | 0.27 | 12.9 | 0.001 | 0.012 | 0.010 | 0.013 | 0.013 | 0.1 | 0.751 | 23.7 | 10.4 | 17.4 | 11.0 | 7.4 | 0.008 |
| Precentral Gyrus L | 0.48 | 0.08 | 0.48 | 0.10 | 0.0 | 0.000 | 0.71 | 0.20 | 0.61 | 0.25 | 3.3 | 0.074 | 0.017 | 0.018 | 0.017 | 0.016 | 0.0 | 0.957 | 22.1 | 12.3 | 16.9 | 11.5 | 4.1 | 0.047 |
| Precentral Gyrus R | 0.49 | 0.08 | 0.49 | 0.11 | 0.0 | 0.000 | 0.77 | 0.10 | 0.60 | 0.23 | 16.1 | 0.000 | 0.017 | 0.017 | 0.022 | 0.022 | 1.4 | 0.247 | 22.2 | 11.0 | 19.1 | 13.2 | 1.3 | 0.249 |
| Postcentral Gyrus L | 0.45 | 0.12 | 0.49 | 0.11 | 3.0 | 0.000 | 0.75 | 0.23 | 0.63 | 0.23 | 5.1 | 0.027 | 0.013 | 0.017 | 0.017 | 0.016 | 1.9 | 0.170 | 18.9 | 12.6 | 18.3 | 10.6 | 0.0 | 0.834 |
| Lateral Occipital Cortex, Superior Division L | 0.52 | 0.07 | 0.54 | 0.10 | 0.5 | 0.000 | 0.75 | 0.09 | 0.65 | 0.16 | 8.2 | 0.006 | 0.028 | 0.029 | 0.027 | 0.019 | 0.0 | 0.874 | 28.5 | 10.0 | 25.2 | 10.0 | 2.3 | 0.136 |
| Lateral Occipital Cortex, Superior Division R | 0.49 | 0.13 | 0.56 | 0.07 | 11.8 | 0.000 | 0.75 | 0.08 | 0.65 | 0.10 | 20.3 | 0.000 | 0.020 | 0.020 | 0.039 | 0.029 | 12.2 | 0.001 | 25.2 | 10.9 | 28.2 | 10.8 | 1.6 | 0.206 |
| Occipital Pole R | 0.43 | 0.14 | 0.50 | 0.12 | 5.6 | 0.000 | 0.57 | 0.26 | 0.61 | 0.19 | 0.5 | 0.494 | 0.021 | 0.028 | 0.027 | 0.030 | 1.0 | 0.331 | 18.0 | 13.5 | 19.8 | 13.1 | 0.4 | 0.531 |
| Cingulate Gyrus, Poster Division | 0.51 | 0.11 | 0.55 | 0.10 | 3.5 | 0.000 | 0.79 | 0.09 | 0.69 | 0.07 | 28.1 | 0.000 | 0.017 | 0.018 | 0.030 | 0.019 | 10.5 | 0.002 | 27.6 | 9.9 | 27.6 | 9.6 | 0.0 | 0.997 |
| Precuneus Cortex | 0.57 | 0.12 | 0.62 | 0.07 | 6.4 | 0.000 | 0.70 | 0.07 | 0.64 | 0.06 | 15.8 | 0.000 | 0.053 | 0.033 | 0.070 | 0.040 | 4.2 | 0.043 | 37.8 | 11.4 | 38.2 | 9.3 | 0.0 | 0.853 |
| Top 30% of Edges |  |  |  |  |  |  |  |  |  |  |  |  |  |  |  |  |  |  |  |  |  |  |  |  |
| Frontal Pole L | 0.82 | 0.04 | 0.79 | 0.05 | 11.1 | 0.001 | 0.75 | 0.03 | 0.74 | 0.04 | 3.6 | 0.062 | 0.0265 | 0.0127 | 0.0207 | 0.0092 | 6.0 | 0.016 | 68.6 | 8.8 | 61.8 | 10.5 | 10.7 | 0.002 |
| Frontal Pole R | 0.84 | 0.03 | 0.81 | 0.05 | 12.8 | 0.001 | 0.74 | 0.02 | 0.73 | 0.03 | 1.8 | 0.178 | 0.0303 | 0.0117 | 0.0221 | 0.0073 | 15.9 | 0.000 | 71.8 | 6.8 | 65.4 | 9.0 | 13.1 | 0.000 |
| Middle Frontal Gyrus L | 0.76 | 0.04 | 0.72 | 0.06 | 11.1 | 0.001 | 0.78 | 0.03 | 0.75 | 0.05 | 14.4 | 0.000 | 0.0145 | 0.0059 | 0.0121 | 0.0065 | 3.2 | 0.078 | 55.5 | 8.6 | 48.0 | 12.2 | 10.4 | 0.002 |
| Middle Frontal Gyrus R | 0.78 | 0.07 | 0.75 | 0.07 | 6.4 | 0.014 | 0.77 | 0.05 | 0.74 | 0.05 | 10.4 | 0.002 | 0.0182 | 0.0071 | 0.0143 | 0.0069 | 6.5 | 0.012 | 60.6 | 11.4 | 52.5 | 15.0 | 7.7 | 0.007 |
| Superior frontal Gyrus R | 0.72 | 0.08 | 0.69 | 0.07 | 4.9 | 0.029 | 0.77 | 0.13 | 0.74 | 0.10 | 2.5 | 0.117 | 0.0108 | 0.0059 | 0.0083 | 0.0045 | 4.7 | 0.033 | 48.3 | 13.6 | 40.9 | 14.4 | 5.9 | 0.017 |
| Inferior Frontal Gyrus, Pars Triangular R | 0.70 | 0.05 | 0.69 | 0.05 | 0.4 | 0.554 | 0.82 | 0.07 | 0.76 | 0.06 | 16.9 | 0.000 | 0.0068 | 0.0053 | 0.0086 | 0.0047 | 2.9 | 0.095 | 42.0 | 10.3 | 40.8 | 11.5 | 0.3 | 0.614 |
| Frontal Orbital Cortex R | 0.72 | 0.07 | 0.69 | 0.06 | 3.0 | 0.087 | 0.78 | 0.07 | 0.75 | 0.08 | 4.8 | 0.031 | 0.0107 | 0.0056 | 0.0088 | 0.0047 | 2.9 | 0.095 | 46.6 | 14.0 | 41.5 | 11.7 | 3.4 | 0.067 |
| Precentral Gyrus L | 0.72 | 0.08 | 0.69 | 0.07 | 2.4 | 0.123 | 0.78 | 0.07 | 0.73 | 0.08 | 11.4 | 0.001 | 0.0109 | 0.0064 | 0.0105 | 0.0077 | 0.1 | 0.796 | 46.3 | 16.5 | 40.9 | 15.8 | 2.4 | 0.128 |
| Precentral Gyrus R | 0.73 | 0.06 | 0.70 | 0.08 | 3.6 | 0.061 | 0.79 | 0.05 | 0.73 | 0.07 | 20.4 | 0.000 | 0.0125 | 0.0070 | 0.0110 | 0.0070 | 1.1 | 0.308 | 48.6 | 12.2 | 42.9 | 15.9 | 3.4 | 0.070 |
| Postcentral Gyrus L | 0.70 | 0.08 | 0.70 | 0.08 | 0.0 | 0.835 | 0.78 | 0.09 | 0.72 | 0.12 | 6.9 | 0.010 | 0.0094 | 0.0055 | 0.0121 | 0.0081 | 3.3 | 0.074 | 43.4 | 15.6 | 44.3 | 15.8 | 0.1 | 0.787 |
| Lateral Occipital Cortex, Superior Division L | 0.77 | 0.05 | 0.75 | 0.06 | 3.2 | 0.078 | 0.77 | 0.03 | 0.74 | 0.04 | 21.3 | 0.000 | 0.0173 | 0.0086 | 0.0150 | 0.0075 | 1.8 | 0.181 | 56.9 | 10.2 | 52.8 | 12.3 | 2.8 | 0.098 |
| Lateral Occipital Cortex, Superior Division R | 0.75 | 0.06 | 0.76 | 0.06 | 0.1 | 0.721 | 0.77 | 0.04 | 0.75 | 0.04 | 7.3 | 0.008 | 0.0142 | 0.0073 | 0.0159 | 0.0080 | 0.9 | 0.336 | 53.7 | 13.0 | 54.9 | 11.3 | 0.2 | 0.651 |
| Occipital Pole R | 0.71 | 0.09 | 0.71 | 0.07 | 0.0 | 0.924 | 0.75 | 0.08 | 0.74 | 0.07 | 0.5 | 0.502 | 0.0137 | 0.0105 | 0.0110 | 0.0070 | 2.0 | 0.163 | 44.7 | 17.5 | 44.4 | 14.8 | 0.0 | 0.935 |
| Cingulate Gyrus, Poster Division | 0.76 | 0.07 | 0.76 | 0.07 | 0.0 | 0.991 | 0.77 | 0.13 | 0.75 | 0.06 | 0.4 | 0.554 | 0.0158 | 0.0104 | 0.0148 | 0.0069 | 0.3 | 0.584 | 54.9 | 12.8 | 54.8 | 10.7 | 0.0 | 0.964 |
| Precuneus Cortex | 0.80 | 0.06 | 0.80 | 0.05 | 0.2 | 0.682 | 0.76 | 0.03 | 0.73 | 0.04 | 16.4 | 0.000 | 0.0208 | 0.0098 | 0.0219 | 0.0086 | 0.3 | 0.581 | 62.7 | 12.3 | 63.9 | 8.9 | 0.3 | 0.589 |

<sup>1</sup>Local efficiency could not be calculated for some regions for up to 7 younger and 5 older participants at the top 10% of edges where the node had no defined local neighbours.

### 2.4 Age Differences in the Whole Brain Metabolic Function

The topology of the age differences in all of the metabolic regions across the whole brain largely followed those for the hubs. Older adults showed significantly higher global efficiency across the whole brain at the top 10% of edges ($t(84) = 3.8$, $p < .01$) but at the top 30% the age difference was not significant ($t(84) = 0.3$, $p = .772$). Older adults had higher global efficiency at the top 10% of edges primarily in regions in the temporal and occipital lobes and the subcortex. At the top 30% of edges, the direction of age group differences reversed, with regions in the temporal and occipital lobes and subcortex showing higher global efficiency in younger adults. Regions in the frontal cortex also showed higher efficiency in younger adults at the top 30% of edges.

Younger adults showed significantly higher local efficiency than older adults across the whole brain at the top 10% of edges ($t(84) = 4.0$, $p < .01$) but the age difference at the top 30% of edges was not significant ($t(84) = 1.9$, $p = .059$). Regions with higher local efficiency in younger adults were seen across most of the cortex, particularly in the frontal, temporal and parietal lobes at the top 30% of edges (Figure S4).

Younger adults had higher betweenness centrality at the whole brain level at both the top 10% of edges ($t(84) = 2.7$, $p = .009$) and top 30% of edges ($t(84) = 2.7$, $p = .009$). At the top 10% of edges, none of the regional age differences survived FDR correction. However, at the top 30% of edges, younger adults had significantly higher betweenness centrality in regions in the frontal and temporal cortices and subcortex (Figure S4).

As the graphs were thresholded based on cost to obtain the top 10% and 30% of edges, the same number of edges were present for older and younger adults across the entire graph and hence whole brain age differences in degree were not tested. At the region level, younger adults showed higher degree in the frontal lobe regions but lower degree in regions in the motor and temporal cortices and subcortex (Figure S4).

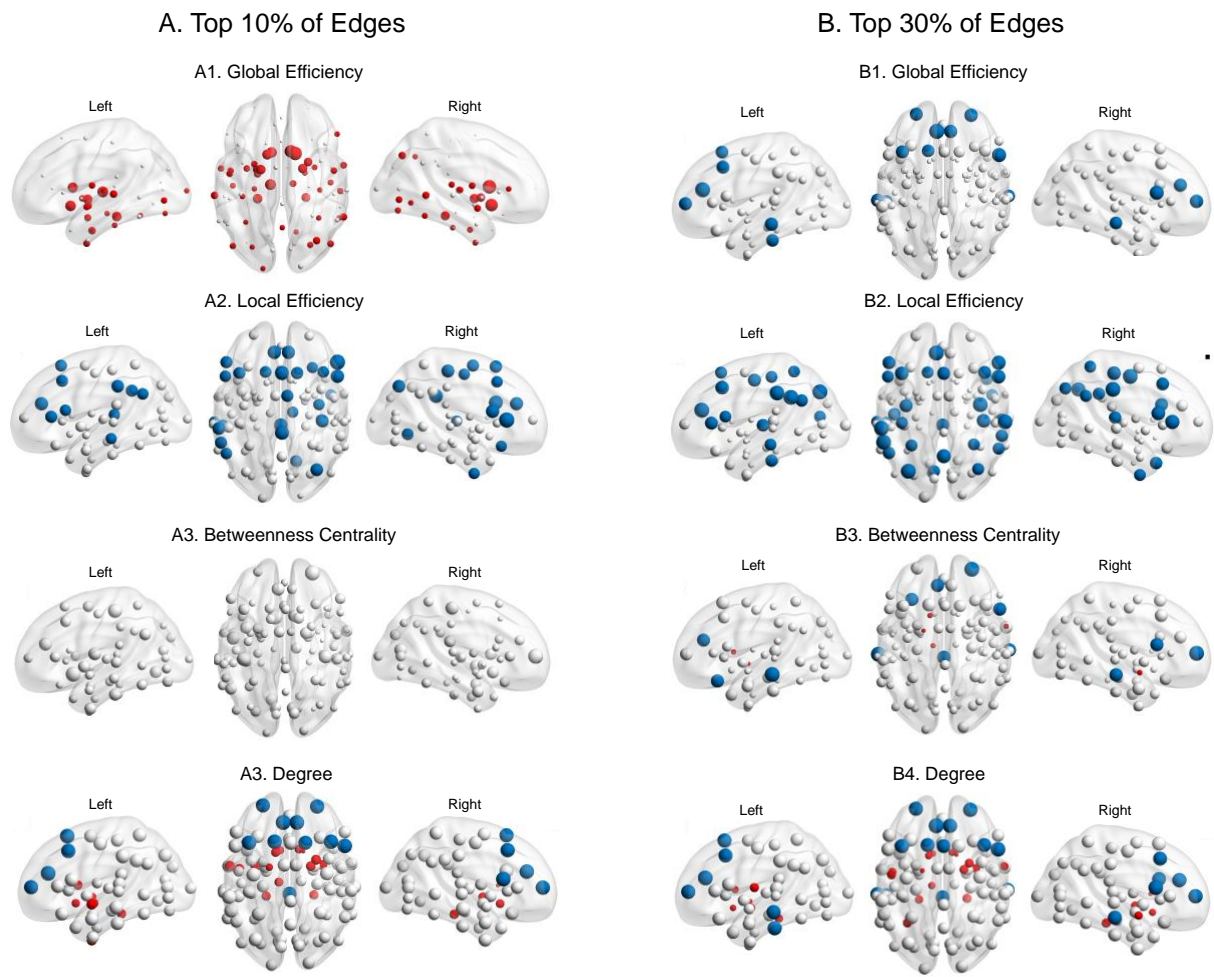

**Figure S4.** Graph metrics results for *all nodes* of the metabolic network for younger and older adults at top 10% (A) and 30% (B) of network edges. Nodes showing age group differences are coloured red for older > younger and blue for younger > older ($p\text{-FDR} < .05$). Nodes showing non-significant age group differences are shaded grey. Size of the node reflects the relative magnitude of the T-statistic (see Table S2 and S3).

**Table S2.** Mean, Standard Deviation (SD) and *t*-test (df = 84) of age group differences in graph metrics at top 10% of network edges.

|  | Global Efficiency |  |  |  |  |  | Local Efficiency |  |  |  |  |  | Betweenness Centrality |  |  |  |  |  | Degree |  |  |  |  |  |
| --- | --- | --- | --- | --- | --- | --- | --- | --- | --- | --- | --- | --- | --- | --- | --- | --- | --- | --- | --- | --- | --- | --- | --- | --- |
|  | Younger |  | Older |  | Young vs Old |  | Younger |  | Older |  | Young vs Old |  | Younger |  | Older |  | Young vs Old |  | Younger |  | Older |  | Young vs Old |  |
|  | Mean | SD | Mean | SD | T | p FDR | Mean | SD | Mean | SD | T | p FDR | Mean | SD | Mean | SD | T | p FDR | Mean | SD | Mean | SD | T | p FDR |
| Frontal Orbital Cortex L | 0.44 | 0.09 | 0.45 | 0.12 | -0.32 | 0.827 | 0.72 | 0.23 | 0.59 | 0.30 | 2.1 | 0.111 | 0.013 | 0.017 | 0.068 | 0.037 | -1.9 | 0.325 | 15.5 | 10.0 | 13.0 | 8.6 | 1.2 | 0.362 |
| Frontal Pole L | 0.60 | 0.06 | 0.61 | 0.07 | -0.68 | 0.622 | 0.69 | 0.06 | 0.66 | 0.09 | 1.7 | 0.205 | 0.004 | 0.008 | 0.004 | 0.008 | -0.7 | 0.696 | 5.7 | 5.9 | 37.1 | 10.6 | 2.8 | 0.027 |
| Frontal Operculum Cortex L | 0.32 | 0.15 | 0.37 | 0.16 | -1.64 | 0.181 | 0.63 | 0.35 | 0.55 | 0.32 | 0.9 | 0.540 | 0.025 | 0.027 | 0.025 | 0.027 | -2.2 | 0.272 | 25.5 | 12.5 | 7.3 | 6.1 | -1.2 | 0.364 |
| Superior Frontal Gyrus L | 0.50 | 0.11 | 0.47 | 0.12 | 1.20 | 0.348 | 0.77 | 0.07 | 0.66 | 0.20 | 3.2 | 0.011 | 0.022 | 0.025 | 0.022 | 0.025 | -0.4 | 0.862 | 29.1 | 10.3 | 16.4 | 11.3 | 3.6 | 0.006 |
| Middle Frontal Gyrus L | 0.53 | 0.06 | 0.52 | 0.07 | 0.10 | 0.964 | 0.77 | 0.07 | 0.68 | 0.14 | 3.3 | 0.011 | 0.007 | 0.009 | 0.007 | 0.009 | 0.2 | 0.962 | 16.0 | 9.6 | 21.8 | 10.1 | 3.3 | 0.010 |
| Inferior Frontal Gyrus; Pars Triangularis L | 0.44 | 0.09 | 0.46 | 0.10 | -1.14 | 0.369 | 0.81 | 0.15 | 0.64 | 0.27 | 3.4 | 0.009 | 0.005 | 0.008 | 0.005 | 0.008 | -0.1 | 0.966 | 13.1 | 8.6 | 13.5 | 7.5 | 1.4 | 0.306 |
| Inferior Frontal Gyrus; pars opercularis L | 0.42 | 0.10 | 0.42 | 0.13 | 0.03 | 0.986 | 0.80 | 0.20 | 0.63 | 0.30 | 2.9 | 0.023 | 0.010 | 0.014 | 0.010 | 0.014 | -2.5 | 0.222 | 18.4 | 7.9 | 9.9 | 7.2 | 1.9 | 0.146 |
| Paracingulate Gyrus L | 0.47 | 0.06 | 0.44 | 0.10 | 1.65 | 0.179 | 0.84 | 0.08 | 0.63 | 0.29 | 4.3 | 0.001 | 0.001 | 0.004 | 0.001 | 0.004 | 1.3 | 0.493 | 2.6 | 2.9 | 11.0 | 7.4 | 4.5 | 0.001 |
| Insular Cortex L | 0.25 | 0.16 | 0.30 | 0.18 | -1.44 | 0.252 | 0.68 | 0.39 | 0.42 | 0.36 | 2.3 | 0.090 | 0.009 | 0.022 | 0.009 | 0.022 | 0.7 | 0.715 | 3.2 | 5.1 | 3.9 | 3.9 | -1.7 | 0.197 |
| Amygdala I | 0.20 | 0.16 | 0.31 | 0.14 | -3.33 | 0.009 | 0.29 | 0.38 | 0.21 | 0.33 | 0.9 | 0.547 | 0.003 | 0.005 | 0.003 | 0.005 | -0.7 | 0.699 | 8.2 | 7.7 | 4.2 | 3.7 | -1.1 | 0.442 |
| Juxtapositional Lobule Cortex - L | 0.31 | 0.18 | 0.38 | 0.13 | -2.09 | 0.088 | 0.75 | 0.34 | 0.57 | 0.34 | 2.2 | 0.105 | 0.017 | 0.018 | 0.017 | 0.018 | 0.2 | 0.975 | 22.1 | 12.3 | 7.0 | 6.6 | 0.8 | 0.581 |
| Precentral Gyrus L | 0.48 | 0.08 | 0.48 | 0.10 | -0.08 | 0.967 | 0.71 | 0.23 | 0.64 | 0.26 | 1.5 | 0.265 | 0.013 | 0.017 | 0.013 | 0.017 | -0.9 | 0.691 | 18.9 | 12.6 | 16.9 | 11.5 | 2.0 | 0.125 |
| PostCG I Postcentral Gyrus L | 0.45 | 0.12 | 0.49 | 0.11 | -1.73 | 0.160 | 0.76 | 0.22 | 0.65 | 0.21 | 2.2 | 0.101 | 0.003 | 0.009 | 0.003 | 0.009 | 1.9 | 0.324 | 4.1 | 5.3 | 18.3 | 10.6 | 0.2 | 0.867 |
| Central Opercular Cortex L | 0.27 | 0.17 | 0.37 | 0.15 | -3.03 | 0.016 | 0.51 | 0.38 | 0.50 | 0.34 | 0.1 | 0.925 | 0.007 | 0.011 | 0.007 | 0.011 | 1.2 | 0.518 | 15.0 | 10.4 | 6.9 | 5.4 | -2.4 | 0.066 |
| Superior Parietal Lobule L | 0.41 | 0.14 | 0.42 | 0.14 | -0.38 | 0.816 | 0.79 | 0.19 | 0.68 | 0.25 | 2.1 | 0.114 | 0.005 | 0.009 | 0.005 | 0.009 | -0.8 | 0.711 | 10.6 | 9.0 | 11.1 | 7.7 | -2.0 | 0.136 |
| Supramarginal Gyrus; Anterior Division L | 0.39 | 0.13 | 0.39 | 0.11 | 0.00 | 0.997 | 0.79 | 0.25 | 0.50 | 0.39 | 3.8 | 0.003 | 0.007 | 0.010 | 0.007 | 0.010 | -1.0 | 0.626 | 15.2 | 9.3 | 6.9 | 5.8 | 2.3 | 0.073 |
| Supramarginal Gyrus; Posterior Division L | 0.43 | 0.10 | 0.44 | 0.12 | -0.33 | 0.828 | 0.75 | 0.23 | 0.56 | 0.27 | 3.4 | 0.008 | 0.004 | 0.008 | 0.004 | 0.008 | -1.1 | 0.548 | 10.0 | 8.2 | 12.1 | 7.5 | -1.7 | 0.208 |
| Angular Gyrus L | 0.40 | 0.10 | 0.44 | 0.10 | -1.79 | 0.144 | 0.77 | 0.24 | 0.59 | 0.29 | 3.0 | 0.020 | 0.005 | 0.009 | 0.005 | 0.009 | 0.5 | 0.799 | 8.3 | 7.7 | 10.7 | 6.9 | -0.4 | 0.780 |
| Parietal Operculum Cortex L | 0.34 | 0.16 | 0.40 | 0.14 | -1.71 | 0.163 | 0.69 | 0.29 | 0.51 | 0.31 | 2.4 | 0.083 | 0.007 | 0.012 | 0.007 | 0.012 | -1.2 | 0.533 | 2.4 | 3.2 | 7.8 | 5.9 | 0.3 | 0.821 |
| Parahippocampal Gyrus; Anterior Division L | 0.18 | 0.17 | 0.30 | 0.15 | -3.52 | 0.007 | 0.20 | 0.26 | 0.17 | 0.22 | 0.5 | 0.755 | 0.003 | 0.012 | 0.003 | 0.012 | -1.4 | 0.465 | 0.8 | 1.0 | 3.8 | 3.0 | -2.1 | 0.115 |
| Parahippocampal Gyrus; Posterior Division L | 0.12 | 0.15 | 0.27 | 0.16 | -4.47 | 0.000 | 0.23 | 0.29 | 0.25 | 0.35 | -0.2 | 0.919 | 0.010 | 0.020 | 0.010 | 0.020 | -1.8 | 0.360 | 3.5 | 4.4 | 3.0 | 3.8 | -3.5 | 0.006 |
| Hippocampus I | 0.22 | 0.17 | 0.30 | 0.15 | -2.51 | 0.038 | 0.35 | 0.37 | 0.28 | 0.35 | 0.7 | 0.671 | 0.002 | 0.008 | 0.002 | 0.008 | -0.9 | 0.698 | 1.4 | 2.6 | 4.0 | 4.1 | -0.6 | 0.670 |
| Thalamus L | 0.13 | 0.18 | 0.33 | 0.17 | -5.07 | 0.000 | 0.51 | 0.46 | 0.42 | 0.37 | 0.6 | 0.692 | 0.005 | 0.010 | 0.005 | 0.010 | -2.3 | 0.232 | 3.4 | 4.9 | 4.8 | 5.0 | -3.8 | 0.004 |
| Caudate L | 0.21 | 0.18 | 0.38 | 0.16 | -4.68 | 0.000 | 0.38 | 0.36 | 0.44 | 0.31 | -0.6 | 0.709 | 0.017 | 0.018 | 0.017 | 0.018 | -0.6 | 0.701 | 7.6 | 6.5 | 8.1 | 6.9 | -3.6 | 0.006 |
| Putamen L | 0.33 | 0.16 | 0.43 | 0.14 | -3.05 | 0.015 | 0.48 | 0.32 | 0.51 | 0.27 | -0.6 | 0.707 | 0.004 | 0.010 | 0.004 | 0.010 | -1.1 | 0.967 | 2.0 | 1.4 | 11.6 | 9.2 | -2.3 | 0.075 |
| Pallidum L | 0.24 | 0.14 | 0.36 | 0.11 | -4.44 | 0.000 | 0.27 | 0.36 | 0.30 | 0.32 | -0.4 | 0.793 | 0.004 | 0.008 | 0.004 | 0.008 | -1.6 | 0.436 | 2.2 | 2.1 | 4.9 | 3.9 | -4.6 | 0.001 |
| Accumbens L | 0.18 | 0.16 | 0.34 | 0.14 | -4.89 | 0.000 | 0.35 | 0.42 | 0.38 | 0.30 | -0.2 | 0.896 | 0.003 | 0.006 | 0.003 | 0.006 | 2.4 | 0.203 | 4.3 | 6.3 | 4.9 | 3.7 | -4.1 | 0.003 |
| Temporal Pole L | 0.26 | 0.17 | 0.31 | 0.18 | -1.38 | 0.267 | 0.47 | 0.41 | 0.36 | 0.35 | 1.0 | 0.489 | 0.003 | 0.009 | 0.003 | 0.009 | -1.2 | 0.533 | 1.5 | 1.8 | 4.5 | 4.9 | -0.2 | 0.865 |
| Planum Polare L | 0.19 | 0.16 | 0.30 | 0.17 | -2.97 | 0.017 | 0.22 | 0.39 | 0.40 | 0.35 | -1.6 | 0.260 | 0.003 | 0.009 | 0.003 | 0.009 | -2.0 | 0.289 | 2.0 | 2.6 | 3.6 | 3.2 | -3.7 | 0.005 |
| Superior Temporal Gyrus; Anterior Division L | 0.20 | 0.15 | 0.30 | 0.15 | -3.02 | 0.015 | 0.33 | 0.40 | 0.29 | 0.34 | 0.3 | 0.816 | 0.002 | 0.004 | 0.002 | 0.004 | -1.0 | 0.617 | 5.2 | 7.1 | 3.6 | 3.2 | -2.6 | 0.048 |
| Superior Temporal Gyrus; Posterior Division L | 0.31 | 0.14 | 0.40 | 0.11 | -3.50 | 0.007 | 0.58 | 0.40 | 0.45 | 0.33 | 1.4 | 0.289 | 0.010 | 0.011 | 0.010 | 0.011 | 0.1 | 0.976 | 4.9 | 6.5 | 7.4 | 6.1 | -1.6 | 0.234 |
| Middle Temporal Gyrus; Anterior Division L | 0.29 | 0.16 | 0.31 | 0.15 | -0.64 | 0.649 | 0.34 | 0.33 | 0.18 | 0.25 | 2.2 | 0.099 | 0.016 | 0.017 | 0.016 | 0.017 | -0.8 | 0.690 | 17.7 | 11.2 | 4.0 | 3.6 | 0.8 | 0.579 |
| Middle Temporal Gyrus; Posterior Division L | 0.45 | 0.10 | 0.44 | 0.13 | 0.48 | 0.739 | 0.66 | 0.23 | 0.50 | 0.29 | 2.7 | 0.037 | 0.003 | 0.006 | 0.003 | 0.006 | -0.2 | 0.984 | 2.6 | 3.2 | 12.8 | 9.0 | 2.2 | 0.079 |
| Inferior Temporal Gyrus; Anterior Division L | 0.21 | 0.17 | 0.28 | 0.17 | -1.92 | 0.122 | 0.45 | 0.39 | 0.24 | 0.35 | 1.9 | 0.165 | 0.010 | 0.014 | 0.010 | 0.014 | 0.7 | 0.697 | 9.5 | 7.3 | 3.0 | 3.0 | -0.6 | 0.674 |
| Inferior Temporal Gyrus; Posterior Division L | 0.39 | 0.11 | 0.39 | 0.15 | 0.20 | 0.892 | 0.60 | 0.33 | 0.47 | 0.30 | 1.7 | 0.219 | 0.013 | 0.015 | 0.013 | 0.015 | 0.8 | 0.693 | 14.5 | 7.3 | 7.9 | 7.4 | 1.0 | 0.460 |
| Planum Temporale L | 0.43 | 0.10 | 0.46 | 0.12 | -1.39 | 0.265 | 0.72 | 0.24 | 0.56 | 0.27 | 2.8 | 0.034 | 0.008 | 0.013 | 0.008 | 0.013 | -1.5 | 0.465 | 7.5 | 6.3 | 13.9 | 8.5 | 0.4 | 0.798 |
| Heschl's Gyrus L | 0.37 | 0.12 | 0.44 | 0.11 | -2.78 | 0.024 | 0.63 | 0.34 | 0.54 | 0.27 | 1.4 | 0.315 | 0.004 | 0.010 | 0.004 | 0.010 | -0.7 | 0.717 | 2.0 | 1.8 | 10.5 | 7.2 | -2.1 | 0.118 |
| Temporal Fusiform Cortex; Anterior Division L | 0.20 | 0.16 | 0.31 | 0.14 | -3.13 | 0.014 | 0.33 | 0.37 | 0.19 | 0.28 | 1.6 | 0.259 | 0.007 | 0.014 | 0.007 | 0.014 | -1.4 | 0.453 | 3.0 | 3.0 | 4.0 | 3.9 | -3.0 | 0.020 |
| Temporal Fusiform Cortex; Posterior Division L | 0.23 | 0.18 | 0.33 | 0.17 | -2.62 | 0.033 | 0.36 | 0.35 | 0.30 | 0.25 | 0.7 | 0.675 | 0.004 | 0.012 | 0.004 | 0.012 | 0.6 | 0.728 | 2.8 | 3.4 | 5.0 | 4.1 | -2.6 | 0.052 |
| Temporal Occipital Fusiform Cortex L | 0.26 | 0.15 | 0.34 | 0.15 | -2.37 | 0.050 | 0.46 | 0.42 | 0.41 | 0.36 | 0.5 | 0.759 | 0.006 | 0.009 | 0.006 | 0.009 | -0.8 | 0.701 | 7.7 | 6.8 | 4.7 | 3.7 | -2.4 | 0.061 |
| Middle Temporal Gyrus; Temporooccipital Part L | 0.34 | 0.15 | 0.40 | 0.14 | -1.91 | 0.121 | 0.60 | 0.37 | 0.55 | 0.26 | 0.7 | 0.677 | 0.005 | 0.012 | 0.005 | 0.012 | -0.9 | 0.686 | 4.6 | 4.9 | 8.8 | 7.1 | -0.7 | 0.588 |
| Inferior Temporal Gyrus; Temporooccipital Part L | 0.29 | 0.16 | 0.36 | 0.15 | -2.22 | 0.069 | 0.57 | 0.41 | 0.43 | 0.36 | 1.6 | 0.255 | 0.008 | 0.011 | 0.008 | 0.011 | -0.1 | 0.959 | 8.0 | 7.5 | 5.4 | 4.4 | -0.8 | 0.543 |
| Occipital Fusiform Gyrus L | 0.34 | 0.15 | 0.42 | 0.14 | -2.46 | 0.043 | 0.57 | 0.33 | 0.54 | 0.34 | 0.3 | 0.866 | 0.006 | 0.009 | 0.006 | 0.009 | 0.0 | 0.969 | 6.1 | 4.2 | 11.3 | 8.2 | -1.9 | 0.145 |
| Supracalcarine Cortex L | 0.36 | 0.08 | 0.39 | 0.13 | -1.12 | 0.374 | 0.50 | 0.35 | 0.41 | 0.32 | 1.2 | 0.386 | 0.005 | 0.008 | 0.005 | 0.008 | -0.1 | 0.982 | 5.7 | 6.2 | 7.8 | 5.0 | -1.6 | 0.210 |
| Cuneal Cortex L | 0.32 | 0.16 | 0.39 | 0.12 | -2.18 | 0.073 | 0.67 | 0.30 | 0.51 | 0.35 | 2.0 | 0.145 | 0.004 | 0.010 | 0.004 | 0.010 | -0.7 | 0.711 | 6.1 | 6.3 | 7.0 | 5.1 | -1.0 | 0.458 |
| Lingual Gyrus L | 0.33 | 0.15 | 0.37 | 0.17 | -1.41 | 0.261 | 0.63 |  |  |  |  |  |  |  |  |  |  |  |  |  |  |  |  |  |

Table S2 continued

|  | Global Efficiency |  |  |  |  |  | Local Efficiency |  |  |  |  |  | Betweenness Centrality |  |  |  |  |  | Degree |  |  |  |  |  |
| --- | --- | --- | --- | --- | --- | --- | --- | --- | --- | --- | --- | --- | --- | --- | --- | --- | --- | --- | --- | --- | --- | --- | --- | --- |
|  | Younger |  | Older |  | Young vs Old |  | Younger |  | Older |  | Young vs Old |  | Younger |  | Older |  | Young vs Old |  | Younger |  | Older |  | Young vs Old |  |
|  | Mean | SD | Mean | SD | T | p FDR | Mean | SD | Mean | SD | T | p FDR | Mean | SD | Mean | SD | T | p FDR | Mean | SD | Mean | SD | T | p FDR |
| Frontal Orbital Cortex R | 0.43 | 0.15 | 0.46 | 0.12 | -1.00 | 0.434 | 0.78 | 0.16 | 0.66 | 0.23 | 2.7 | 0.037 | 0.098 | 0.049 | 0.098 | 0.049 | -0.2 | 0.941 | 48.9 | 8.4 | 14.6 | 7.9 | 0.9 | 0.473 |
| Frontal Pole R | 0.63 | 0.07 | 0.64 | 0.06 | -0.48 | 0.754 | 0.66 | 0.06 | 0.63 | 0.08 | 1.5 | 0.247 | 0.006 | 0.009 | 0.006 | 0.009 | 2.4 | 0.220 | 9.3 | 7.5 | 42.3 | 10.2 | 3.2 | 0.011 |
| Frontal Operculum Cortex R | 0.39 | 0.12 | 0.43 | 0.12 | -1.82 | 0.140 | 0.68 | 0.27 | 0.64 | 0.26 | 0.7 | 0.677 | 0.012 | 0.010 | 0.012 | 0.010 | -2.1 | 0.267 | 23.7 | 10.4 | 10.9 | 6.6 | -1.1 | 0.422 |
| Superior Frontal Gyrus R | 0.49 | 0.09 | 0.48 | 0.08 | 0.35 | 0.817 | 0.81 | 0.08 | 0.66 | 0.26 | 3.6 | 0.005 | 0.477 | 0.073 | 0.477 | 0.073 | -1.3 | 0.461 | 2.2 | 0.3 | 17.4 | 11.0 | 2.7 | 0.037 |
| Middle Frontal Gyrus R | 0.50 | 0.24 | 20.70 | 7.29 | -0.09 | 0.968 | 14.54 | 8.00 | 0.48 | 0.10 | 3.0 | 0.022 | 0.343 | 0.156 | 0.343 | 0.156 | -2.8 | 0.127 | 2.6 | 0.4 | 0.7 | 0.1 | 3.2 | 0.012 |
| Inferior Frontal Gyrus; Pars Triangularis R | 0.49 | 0.25 | 21.73 | 8.90 | -2.54 | 0.038 | 14.93 | 7.83 | 0.28 | 0.17 | 5.6 | 0.000 | 0.293 | 0.178 | 0.293 | 0.178 | -0.1 | 0.968 | 2.5 | 1.0 | 0.4 | 0.3 | -0.2 | 0.880 |
| Inferior Frontal Gyrus; Pars Opercularis R | 0.30 | 0.28 | 4.28 | 4.60 | 0.36 | 0.819 | 5.22 | 4.31 | 0.26 | 0.18 | 4.3 | 0.001 | 0.348 | 0.165 | 0.348 | 0.165 | -1.7 | 0.375 | 2.6 | 0.5 | 0.4 | 0.3 | 3.7 | 0.004 |
| Paracingulate Gyrus R | 0.21 | 0.25 | 4.45 | 5.17 | 0.22 | 0.894 | 5.09 | 4.99 | 0.32 | 0.16 | 4.2 | 0.001 | 0.487 | 0.114 | 0.487 | 0.114 | -0.3 | 0.916 | 2.2 | 0.5 | 0.8 | 0.3 | 3.8 | 0.005 |
| Insular Cortex R | 0.45 | 0.36 | 7.30 | 8.49 | -1.83 | 0.140 | 6.67 | 7.38 | 0.49 | 0.08 | 1.6 | 0.267 | 0.434 | 0.149 | 0.434 | 0.149 | -3.2 | 0.089 | 2.4 | 0.8 | 0.6 | 0.2 | -1.0 | 0.466 |
| Amygdala r | 0.43 | 0.21 | 22.15 | 11.01 | -0.89 | 0.483 | 19.09 | 13.15 | 0.42 | 0.09 | 1.9 | 0.152 | 0.413 | 0.116 | 0.413 | 0.116 | 0.1 | 0.960 | 2.4 | 0.4 | 0.7 | 0.2 | -0.6 | 0.671 |
| Juxtapositional Lobule Cortex R | 0.37 | 0.23 | 12.83 | 10.01 | -0.90 | 0.478 | 14.54 | 11.56 | 0.32 | 0.17 | 3.4 | 0.010 | 0.402 | 0.116 | 0.402 | 0.116 | -1.4 | 0.463 | 2.5 | 0.6 | 0.5 | 0.3 | 0.4 | 0.798 |
| Precentral Gyrus R | 0.48 | 0.31 | 6.55 | 7.05 | -0.05 | 0.979 | 8.43 | 6.83 | 0.35 | 0.17 | 2.9 | 0.027 | 0.353 | 0.145 | 0.353 | 0.145 | -3.5 | 0.081 | 2.7 | 0.6 | 0.6 | 0.3 | 1.2 | 0.400 |
| Postcentral Gyrus R | 0.48 | 0.29 | 9.65 | 9.58 | -0.52 | 0.728 | 8.20 | 6.65 | 0.32 | 0.14 | 4.0 | 0.002 | 0.430 | 0.116 | 0.430 | 0.116 | 1.4 | 0.459 | 2.3 | 0.4 | 0.5 | 0.4 | -0.7 | 0.582 |
| Central Opercular Cortex R | 0.39 | 0.33 | 6.15 | 6.56 | -3.08 | 0.015 | 5.83 | 5.76 | 0.40 | 0.11 | 0.2 | 0.925 | 0.458 | 0.100 | 0.458 | 0.100 | -1.0 | 0.621 | 2.2 | 0.4 | 0.6 | 0.3 | -1.3 | 0.363 |
| Superior Parietal Lobule R | 0.42 | 0.25 | 10.60 | 8.56 | -1.71 | 0.162 | 11.33 | 7.62 | 0.40 | 0.12 | 2.0 | 0.145 | 0.374 | 0.149 | 0.374 | 0.149 | -0.3 | 0.884 | 2.5 | 0.4 | 0.6 | 0.3 | 0.8 | 0.551 |
| Supramarginal Gyrus; Anterior Division R | 0.44 | 0.28 | 12.08 | 9.79 | -0.93 | 0.467 | 14.04 | 9.27 | 0.33 | 0.12 | 0.9 | 0.573 | 0.285 | 0.157 | 0.285 | 0.157 | -1.4 | 0.473 | 2.9 | 1.0 | 0.4 | 0.3 | 0.2 | 0.857 |
| Supramarginal Gyrus; Posterior Division R | 0.32 | 0.26 | 4.95 | 4.84 | -1.23 | 0.339 | 6.65 | 5.49 | 0.19 | 0.17 | 2.2 | 0.098 | 0.279 | 0.165 | 0.279 | 0.165 | -1.9 | 0.333 | 3.0 | 0.8 | 0.2 | 0.4 | -0.4 | 0.773 |
| Angular Gyrus R | 0.21 | 0.32 | 2.13 | 2.39 | -2.22 | 0.068 | 3.48 | 2.54 | 0.13 | 0.16 | 1.7 | 0.216 | 0.294 | 0.179 | 0.294 | 0.179 | -1.4 | 0.470 | 2.6 | 1.0 | 0.3 | 0.4 | -1.0 | 0.476 |
| Parietal Operculum Cortex R | 0.15 | 0.22 | 1.13 | 1.62 | -1.48 | 0.236 | 3.00 | 3.00 | 0.25 | 0.17 | 0.0 | 0.968 | 0.310 | 0.197 | 0.310 | 0.197 | 0.6 | 0.736 | 2.6 | 0.5 | 0.3 | 0.3 | -1.5 | 0.258 |
| Parahippocampal Gyrus; Anterior Division R | 0.27 | 0.27 | 3.75 | 5.14 | -2.80 | 0.023 | 5.09 | 5.59 | 0.18 | 0.19 | 0.3 | 0.836 | 0.397 | 0.132 | 0.397 | 0.132 | -0.7 | 0.709 | 2.5 | 0.6 | 0.7 | 0.4 | -2.5 | 0.051 |
| Parahippocampal Gyrus; Posterior Division R | 0.36 | 0.33 | 2.30 | 4.32 | -4.20 | 0.001 | 5.43 | 7.16 | 0.19 | 0.17 | 1.6 | 0.251 | 0.442 | 0.140 | 0.442 | 0.140 | -2.1 | 0.273 | 2.3 | 0.4 | 0.2 | 0.3 | -3.5 | 0.006 |
| Hippocampus R | 0.34 | 0.24 | 2.53 | 2.50 | -1.22 | 0.342 | 7.78 | 5.31 | 0.30 | 0.18 | 0.5 | 0.751 | 0.346 | 0.115 | 0.346 | 0.115 | -2.2 | 0.294 | 2.8 | 0.5 | 0.3 | 0.3 | -1.1 | 0.402 |
| Thalamus R | 0.33 | 0.18 | 7.03 | 8.27 | -3.22 | 0.012 | 12.35 | 8.18 | 0.22 | 0.15 | 2.8 | 0.038 | 0.325 | 0.113 | 0.325 | 0.113 | 0.2 | 0.990 | 3.0 | 0.7 | 0.2 | 0.3 | -2.4 | 0.063 |
| Caudate R | 0.23 | 0.30 | 1.80 | 1.98 | -6.27 | 0.000 | 3.78 | 3.26 | 0.16 | 0.15 | -2.2 | 0.108 | 0.342 | 0.184 | 0.342 | 0.184 | 0.4 | 0.840 | 2.5 | 0.5 | 0.4 | 0.4 | -5.7 | 0.000 |
| Putamen R | 0.35 | 0.38 | 1.58 | 1.30 | -3.98 | 0.002 | 3.30 | 2.51 | 0.30 | 0.17 | -0.9 | 0.551 | 0.317 | 0.160 | 0.317 | 0.160 | -0.8 | 0.698 | 2.8 | 0.6 | 0.5 | 0.4 | -3.0 | 0.020 |
| Palidum R | 0.36 | 0.28 | 5.75 | 6.57 | -4.32 | 0.001 | 6.98 | 7.16 | 0.25 | 0.18 | -0.7 | 0.669 | 0.344 | 0.150 | 0.344 | 0.150 | 0.6 | 0.706 | 2.7 | 0.5 | 0.4 | 0.3 | -3.3 | 0.009 |
| Accumbens R | 0.31 | 0.31 | 2.88 | 3.11 | -5.82 | 0.000 | 3.83 | 3.55 | 0.23 | 0.18 | 0.4 | 0.773 | 0.436 | 0.079 | 0.436 | 0.079 | 0.7 | 0.704 | 2.4 | 0.4 | 0.4 | 0.4 | -3.9 | 0.004 |
| Temporal Pole R | 0.33 | 0.28 | 2.60 | 3.06 | -1.08 | 0.394 | 4.39 | 3.41 | 0.39 | 0.09 | 1.3 | 0.341 | 0.390 | 0.110 | 0.390 | 0.110 | -1.2 | 0.509 | 2.6 | 0.5 | 0.5 | 0.3 | -0.8 | 0.546 |
| Planum Polare R | 0.43 | 0.25 | 9.15 | 8.16 | -1.92 | 0.123 | 9.83 | 6.16 | 0.34 | 0.14 | 0.7 | 0.670 | 0.467 | 0.075 | 0.467 | 0.075 | -1.7 | 0.362 | 2.3 | 0.4 | 0.4 | 0.3 | -1.3 | 0.347 |
| Superior Temporal Gyrus; Anterior Division R | 0.29 | 0.23 | 7.08 | 5.71 | -3.17 | 0.013 | 6.52 | 4.34 | 0.46 | 0.09 | 1.0 | 0.508 | 0.344 | 0.141 | 0.344 | 0.141 | -1.7 | 0.365 | 2.8 | 0.8 | 0.5 | 0.2 | -2.5 | 0.052 |
| Superior Temporal Gyrus; Posterior Division R | 0.44 | 0.25 | 18.70 | 9.85 | -2.25 | 0.065 | 14.13 | 8.56 | 0.31 | 0.17 | 0.6 | 0.666 | 0.396 | 0.121 | 0.396 | 0.121 | -2.1 | 0.264 | 2.5 | 0.3 | 0.5 | 0.3 | -0.4 | 0.764 |
| Middle Temporal Gyrus; Anterior Division R | 0.30 | 0.34 | 6.08 | 6.42 | -1.78 | 0.148 | 4.96 | 4.36 | 0.37 | 0.10 | 1.0 | 0.484 | 0.461 | 0.076 | 0.461 | 0.076 | 0.6 | 0.741 | 2.3 | 0.3 | 0.5 | 0.3 | 0.5 | 0.713 |
| Middle Temporal Gyrus; Posterior Division R | 0.36 | 0.29 | 7.63 | 6.25 | -0.22 | 0.888 | 7.24 | 5.35 | 0.41 | 0.08 | 0.1 | 0.926 | 0.411 | 0.122 | 0.411 | 0.122 | -2.4 | 0.197 | 2.5 | 0.5 | 0.5 | 0.2 | 2.3 | 0.074 |
| Inferior Temporal Gyrus; Anterior Division R | 0.44 | 0.22 | 10.73 | 6.94 | -0.98 | 0.441 | 12.46 | 6.79 | 0.33 | 0.15 | 2.8 | 0.028 | 0.286 | 0.159 | 0.286 | 0.159 | -1.5 | 0.469 | 2.9 | 1.0 | 0.5 | 0.3 | 1.0 | 0.471 |
| Inferior Temporal Gyrus; Posterior Division R | 0.46 | 0.27 | 6.35 | 6.10 | -0.94 | 0.464 | 8.67 | 5.84 | 0.19 | 0.16 | 1.6 | 0.262 | 0.326 | 0.147 | 0.326 | 0.147 | -3.2 | 0.057 | 2.8 | 0.9 | 0.1 | 0.3 | 0.3 | 0.820 |
| Planum Temporale R | 0.16 | 0.21 | 2.03 | 2.20 | -2.89 | 0.020 | 3.76 | 3.10 | 0.24 | 0.18 | -0.5 | 0.763 | 0.335 | 0.163 | 0.335 | 0.163 | -0.7 | 0.705 | 2.7 | 0.6 | 0.3 | 0.3 | -1.2 | 0.402 |
| Heschl's Gyrus R | 0.20 | 0.26 | 3.13 | 4.05 | -2.84 | 0.021 | 4.70 | 3.71 | 0.24 | 0.17 | 0.2 | 0.924 | 0.430 | 0.067 | 0.430 | 0.067 | -1.2 | 0.505 | 2.4 | 0.4 | 0.5 | 0.4 | -1.8 | 0.169 |
| Temporal Fusiform Cortex; Anterior Division R | 0.30 | 0.29 | 2.90 | 3.55 | -2.74 | 0.026 | 4.98 | 4.41 | 0.35 | 0.15 | -0.5 | 0.756 | 0.354 | 0.151 | 0.354 | 0.151 | -2.9 | 0.109 | 2.6 | 0.4 | 0.5 | 0.3 | -3.0 | 0.022 |
| Temporal Fusiform Cortex; Posterior Division R | 0.41 | 0.28 | 8.00 | 6.85 | -2.54 | 0.039 | 8.98 | 5.62 | 0.26 | 0.17 | 0.8 | 0.644 | 0.422 | 0.125 | 0.422 | 0.125 | -0.6 | 0.704 | 2.4 | 0.5 | 0.4 | 0.4 | -1.9 | 0.151 |
| Temporal Occipital Fusiform Cortex R | 0.34 | 0.30 | 3.75 | 4.21 | -2.53 | 0.037 | 5.28 | 4.61 | 0.34 | 0.14 | 1.7 | 0.207 | 0.401 | 0.110 | 0.401 | 0.110 | -1.7 | 0.365 | 2.5 | 0.5 | 0.5 | 0.3 | -2.4 | 0.067 |
| Middle Temporal Gyrus; Temporooccipital Part R | 0.43 | 0.32 | 7.08 | 7.35 | -3.43 | 0.007 | 10.37 | 8.96 | 0.39 | 0.09 | 1.9 | 0.150 | 0.401 | 0.133 | 0.401 | 0.133 | -1.0 | 0.625 | 2.5 | 0.5 | 0.4 | 0.3 | -0.7 | 0.578 |
| Inferior Temporal Gyrus; Temporooccipital Part R | 0.32 | 0.20 | 9.50 | 6.48 | -2.74 | 0.025 | 7.74 | 5.20 | 0.36 | 0.11 | 0.7 | 0.668 | 0.396 | 0.133 | 0.396 | 0.133 | -0.5 | 0.798 | 2.6 | 0.5 | 0.5 | 0.3 | -1.6 | 0.227 |
| Occipital Fusiform Gyrus R | 0.45 | 0.25 | 6.93 | 6.59 | -2.91 | 0.019 | 8.30 | 6.15 | 0.32 | 0.16 | 1.1 | 0.428 | 0.410 | 0.111 | 0.410 | 0.111 | -3.2 | 0.069 | 2.6 | 0.6 | 0.6 | 0.3 | -1.8 | 0.157 |
| Supracalcarine Cortex R | 0.34 | 0.30 | 5.65 | 5.13 | -0.48 | 0.747 | 8.50 | 7.63 | 0.36 | 0.14 | 0.7 | 0.665 | 0.561 | 0.068 | 0.561 | 0.068 | 1.1 | 0.573 | 1.9 | 0.2 | 0.5 | 0.3 | 1.4 | 0.309 |
| Cuneal Cortex R | 0.35 | 0.31 | 8.53 | 8.23 | -1.61 | 0.187 | 8.98 | 7.29 | 0.49 | 0.13 | 0.5 | 0.718 | 0.460 | 0.133 | 0.460 | 0.133 | 0.1 | 0.975 | 2.2 | 0.5 | 0.6 | 0.1 | -1.0 | 0.456 |
| Lingual Gyrus R | 0.43 | 0.17 | 25.15 | 10.93 | -2.56 | 0.038 | 28.15 | 10.85 | 0.37 | 0.13 | 3.2 | 0.012 | 0.498 | 0.121 | 0.498 | 0.121 | -1.1 | 0.569 | 2.1 | 0.4 | 0.5 | 0.3 | -2.0 | 0.125 |
| Intracalcarine Cortex R |  |  |  |  |  |  |  |  |  |  |  |  |  |  |  |  |  |  |  |  |  |  |  |  |

**Table S3.** Mean, Standard Deviation (SD) and *t*-test (df = 84) of age group differences in graph metrics at top 30% of network edges.

|  | Global Efficiency |  |  |  |  |  | Local Efficiency |  |  |  |  |  | Betweenness Centrality |  |  |  |  |  | Degree |  |  |  |  |  |
| --- | --- | --- | --- | --- | --- | --- | --- | --- | --- | --- | --- | --- | --- | --- | --- | --- | --- | --- | --- | --- | --- | --- | --- | --- |
|  | Younger |  | Older |  | Young vs Old |  | Younger |  | Older |  | Young vs Old |  | Younger |  | Older |  | Young vs Old |  | Younger |  | Older |  | Young vs Old |  |
|  | Mean | SD | Mean | SD | T | p_FDR | Mean | SD | Mean | SD | T | p_FDR | Mean | SD | Mean | SD | T | p_FDR | Mean | SD | Mean | SD | T | p_FDR |
| Frontal Orbital Cortex L | 0.71 | 0.06 | 0.67 | 0.06 | 2.6 | 0.078 | 0.78 | 0.06 | 0.75 | 0.07 | 1.9 | 0.121 | 0.011 | 0.007 | 0.008 | 0.005 | 2.8 | 0.046 | 44.3 | 13.6 | 37.4 | 13.0 | 2.4 | 0.064 |
| Frontal Pole L | 0.82 | 0.04 | 0.79 | 0.05 | 3.3 | 0.023 | 0.75 | 0.03 | 0.74 | 0.04 | 1.9 | 0.134 | 0.026 | 0.013 | 0.021 | 0.009 | 2.4 | 0.087 | 68.6 | 8.8 | 61.8 | 10.5 | 3.3 | 0.013 |
| Frontal Operculum Cortex L | 0.62 | 0.06 | 0.63 | 0.06 | -0.5 | 0.811 | 0.73 | 0.14 | 0.72 | 0.11 | 0.6 | 0.680 | 0.005 | 0.004 | 0.005 | 0.004 | 0.3 | 0.846 | 27.1 | 12.2 | 28.4 | 12.0 | -0.5 | 0.821 |
| Superior Frontal Gyrus L | 0.73 | 0.08 | 0.68 | 0.08 | 3.1 | 0.040 | 0.78 | 0.03 | 0.74 | 0.10 | 2.7 | 0.037 | 0.013 | 0.008 | 0.008 | 0.006 | 2.9 | 0.069 | 50.2 | 15.5 | 39.3 | 16.1 | 3.2 | 0.013 |
| Middle Frontal Gyrus L | 0.76 | 0.04 | 0.72 | 0.06 | 3.3 | 0.020 | 0.78 | 0.03 | 0.75 | 0.05 | 3.8 | 0.003 | 0.015 | 0.006 | 0.012 | 0.007 | 1.8 | 0.229 | 55.5 | 8.6 | 48.0 | 12.2 | 3.2 | 0.013 |
| Inferior Frontal Gyrus; Pars Triangularis L | 0.70 | 0.06 | 0.68 | 0.05 | 1.6 | 0.287 | 0.80 | 0.07 | 0.75 | 0.08 | 3.3 | 0.009 | 0.008 | 0.005 | 0.008 | 0.004 | 0.0 | 0.982 | 43.1 | 13.1 | 39.2 | 11.1 | 1.5 | 0.297 |
| Inferior Frontal Gyrus; pars opercularis L | 0.67 | 0.07 | 0.64 | 0.06 | 2.1 | 0.148 | 0.79 | 0.08 | 0.75 | 0.08 | 2.6 | 0.041 | 0.007 | 0.005 | 0.006 | 0.005 | 0.7 | 0.725 | 37.9 | 14.0 | 31.5 | 12.8 | 2.2 | 0.099 |
| Paracingulate Gyrus L | 0.71 | 0.05 | 0.65 | 0.06 | 4.7 | 0.001 | 0.81 | 0.04 | 0.74 | 0.09 | 4.7 | 0.001 | 0.010 | 0.007 | 0.006 | 0.004 | 2.8 | 0.045 | 45.5 | 10.5 | 33.4 | 12.4 | 4.8 | 0.001 |
| Insular Cortex L | 0.57 | 0.11 | 0.58 | 0.10 | -0.4 | 0.824 | 0.67 | 0.19 | 0.65 | 0.11 | 0.4 | 0.789 | 0.004 | 0.003 | 0.004 | 0.003 | 0.2 | 0.878 | 20.3 | 10.5 | 21.2 | 9.7 | -0.4 | 0.829 |
| Amygdala l | 0.54 | 0.09 | 0.56 | 0.06 | -1.4 | 0.336 | 0.52 | 0.26 | 0.53 | 0.20 | -0.4 | 0.794 | 0.004 | 0.004 | 0.003 | 0.002 | 1.4 | 0.379 | 15.0 | 11.8 | 16.7 | 8.8 | -0.8 | 0.699 |
| Juxtapositional Lobule Cortex - L | 0.62 | 0.08 | 0.61 | 0.12 | 0.4 | 0.825 | 0.77 | 0.18 | 0.73 | 0.09 | 1.2 | 0.377 | 0.005 | 0.006 | 0.005 | 0.003 | 0.6 | 0.785 | 26.9 | 13.6 | 26.9 | 12.6 | 0.0 | 0.985 |
| Precentral Gyrus L | 0.72 | 0.08 | 0.69 | 0.07 | 1.6 | 0.296 | 0.78 | 0.07 | 0.73 | 0.08 | 3.4 | 0.010 | 0.011 | 0.006 | 0.010 | 0.008 | 0.3 | 0.870 | 46.3 | 16.5 | 40.9 | 15.8 | 1.5 | 0.283 |
| PostCG I Postcentral Gyrus L | 0.70 | 0.08 | 0.70 | 0.08 | -0.2 | 0.885 | 0.78 | 0.09 | 0.72 | 0.12 | 2.6 | 0.042 | 0.009 | 0.006 | 0.012 | 0.008 | -1.8 | 0.230 | 43.4 | 15.6 | 44.3 | 15.8 | -0.3 | 0.887 |
| Central Opercular Cortex L | 0.59 | 0.07 | 0.63 | 0.07 | -2.6 | 0.081 | 0.67 | 0.18 | 0.72 | 0.09 | -1.4 | 0.284 | 0.003 | 0.003 | 0.006 | 0.005 | -2.9 | 0.055 | 20.9 | 12.9 | 28.8 | 12.8 | -2.8 | 0.028 |
| Superior Parietal Lobule L | 0.67 | 0.09 | 0.66 | 0.06 | 1.0 | 0.570 | 0.79 | 0.07 | 0.75 | 0.09 | 2.6 | 0.041 | 0.007 | 0.005 | 0.006 | 0.004 | 1.1 | 0.524 | 37.8 | 16.3 | 34.2 | 12.9 | 1.1 | 0.438 |
| Supramarginal Gyrus; Anterior Division L | 0.66 | 0.07 | 0.63 | 0.06 | 2.4 | 0.089 | 0.81 | 0.08 | 0.73 | 0.09 | 4.5 | 0.000 | 0.005 | 0.004 | 0.005 | 0.004 | 0.5 | 0.821 | 34.9 | 13.8 | 28.2 | 12.1 | 2.4 | 0.062 |
| Supramarginal Gyrus; Posterior Division L | 0.69 | 0.06 | 0.67 | 0.05 | 1.5 | 0.295 | 0.80 | 0.05 | 0.74 | 0.07 | 4.6 | 0.001 | 0.008 | 0.004 | 0.008 | 0.004 | -0.3 | 0.849 | 41.3 | 11.8 | 37.6 | 11.3 | 1.5 | 0.294 |
| Angular Gyrus L | 0.65 | 0.07 | 0.66 | 0.05 | -0.3 | 0.874 | 0.80 | 0.08 | 0.74 | 0.09 | 3.0 | 0.023 | 0.005 | 0.004 | 0.006 | 0.004 | -1.1 | 0.530 | 33.7 | 12.9 | 34.2 | 11.5 | -0.2 | 0.936 |
| Parietal Operculum Cortex L | 0.62 | 0.08 | 0.63 | 0.06 | -0.5 | 0.816 | 0.73 | 0.18 | 0.72 | 0.11 | 0.1 | 0.959 | 0.005 | 0.004 | 0.006 | 0.005 | -1.3 | 0.403 | 28.3 | 14.6 | 29.1 | 10.7 | -0.3 | 0.891 |
| Parahippocampal Gyrus; Anterior Division L | 0.54 | 0.08 | 0.57 | 0.10 | -1.4 | 0.346 | 0.51 | 0.25 | 0.57 | 0.17 | -1.2 | 0.377 | 0.003 | 0.003 | 0.005 | 0.003 | -2.1 | 0.157 | 14.5 | 9.5 | 18.6 | 8.4 | -2.1 | 0.109 |
| Parahippocampal Gyrus; Posterior Division L | 0.52 | 0.06 | 0.56 | 0.11 | -1.7 | 0.236 | 0.53 | 0.28 | 0.57 | 0.19 | -0.7 | 0.613 | 0.002 | 0.002 | 0.003 | 0.003 | -3.1 | 0.054 | 11.2 | 6.5 | 17.0 | 9.7 | -3.2 | 0.012 |
| Hippocampus l | 0.56 | 0.09 | 0.56 | 0.11 | -0.2 | 0.886 | 0.55 | 0.28 | 0.55 | 0.19 | 0.0 | 0.994 | 0.005 | 0.005 | 0.004 | 0.003 | 1.1 | 0.523 | 17.2 | 12.7 | 17.9 | 10.8 | -0.3 | 0.878 |
| Thalamus L | 0.55 | 0.06 | 0.59 | 0.11 | -2.4 | 0.093 | 0.60 | 0.29 | 0.70 | 0.15 | -2.0 | 0.116 | 0.002 | 0.002 | 0.004 | 0.003 | -2.9 | 0.048 | 14.8 | 9.5 | 23.0 | 11.0 | -3.7 | 0.005 |
| Caudate L | 0.58 | 0.07 | 0.63 | 0.11 | -2.5 | 0.075 | 0.64 | 0.22 | 0.71 | 0.07 | -2.0 | 0.123 | 0.004 | 0.004 | 0.007 | 0.004 | -2.8 | 0.046 | 19.7 | 11.5 | 29.7 | 13.1 | -3.7 | 0.005 |
| Putamen L | 0.63 | 0.08 | 0.65 | 0.12 | -0.9 | 0.630 | 0.70 | 0.18 | 0.72 | 0.11 | -0.8 | 0.584 | 0.009 | 0.009 | 0.008 | 0.006 | 0.7 | 0.732 | 29.4 | 14.5 | 34.2 | 14.5 | -1.5 | 0.292 |
| Pallidum L | 0.53 | 0.07 | 0.58 | 0.10 | -2.8 | 0.053 | 0.56 | 0.25 | 0.62 | 0.15 | -1.5 | 0.252 | 0.003 | 0.002 | 0.005 | 0.003 | -3.6 | 0.017 | 12.4 | 7.4 | 21.1 | 10.2 | -4.5 | 0.001 |
| Accumbens L | 0.53 | 0.07 | 0.58 | 0.10 | -2.7 | 0.072 | 0.57 | 0.27 | 0.68 | 0.15 | -2.3 | 0.066 | 0.002 | 0.002 | 0.003 | 0.003 | -1.8 | 0.234 | 13.2 | 9.0 | 21.7 | 9.9 | -4.1 | 0.002 |
| Temporal Pole L | 0.60 | 0.07 | 0.59 | 0.11 | 0.9 | 0.629 | 0.66 | 0.17 | 0.60 | 0.20 | 1.5 | 0.269 | 0.006 | 0.005 | 0.005 | 0.003 | 1.8 | 0.232 | 24.1 | 13.4 | 21.8 | 10.4 | 0.9 | 0.606 |
| Planum Polare L | 0.53 | 0.07 | 0.57 | 0.10 | -1.9 | 0.189 | 0.56 | 0.29 | 0.65 | 0.15 | -1.8 | 0.144 | 0.002 | 0.002 | 0.003 | 0.002 | -2.9 | 0.050 | 12.7 | 8.1 | 19.0 | 9.0 | -3.4 | 0.009 |
| Superior Temporal Gyrus; Anterior Division L | 0.55 | 0.07 | 0.58 | 0.06 | -2.2 | 0.143 | 0.56 | 0.27 | 0.64 | 0.14 | -1.7 | 0.189 | 0.002 | 0.002 | 0.003 | 0.002 | -2.0 | 0.190 | 15.2 | 10.3 | 18.5 | 8.3 | -1.6 | 0.248 |
| Superior Temporal Gyrus; Posterior Division L | 0.60 | 0.07 | 0.63 | 0.05 | -2.3 | 0.109 | 0.73 | 0.17 | 0.73 | 0.09 | 0.1 | 0.976 | 0.004 | 0.004 | 0.005 | 0.004 | -1.9 | 0.196 | 23.4 | 13.9 | 28.7 | 10.7 | -2.0 | 0.134 |
| Middle Temporal Gyrus; Anterior Division L | 0.59 | 0.07 | 0.59 | 0.06 | 0.3 | 0.879 | 0.61 | 0.25 | 0.65 | 0.11 | -1.0 | 0.482 | 0.006 | 0.006 | 0.004 | 0.003 | 1.9 | 0.211 | 22.3 | 12.8 | 20.9 | 9.1 | 0.6 | 0.790 |
| Middle Temporal Gyrus; Posterior Division L | 0.72 | 0.07 | 0.67 | 0.07 | 3.5 | 0.019 | 0.76 | 0.07 | 0.71 | 0.09 | 2.6 | 0.041 | 0.014 | 0.010 | 0.007 | 0.004 | 4.1 | 0.009 | 46.1 | 13.9 | 36.0 | 14.0 | 3.4 | 0.010 |
| Inferior Temporal Gyrus; Anterior Division L | 0.58 | 0.07 | 0.57 | 0.10 | 0.6 | 0.801 | 0.67 | 0.21 | 0.58 | 0.20 | 2.0 | 0.115 | 0.004 | 0.003 | 0.003 | 0.002 | 0.5 | 0.818 | 19.7 | 11.3 | 18.8 | 8.5 | 0.4 | 0.847 |
| Inferior Temporal Gyrus; Posterior Division L | 0.67 | 0.06 | 0.62 | 0.07 | 3.0 | 0.036 | 0.76 | 0.09 | 0.68 | 0.17 | 2.9 | 0.022 | 0.008 | 0.006 | 0.006 | 0.004 | 2.4 | 0.101 | 36.6 | 12.8 | 27.7 | 12.9 | 3.2 | 0.012 |
| Planum Temporale L | 0.70 | 0.06 | 0.68 | 0.05 | 2.0 | 0.165 | 0.79 | 0.06 | 0.75 | 0.09 | 2.4 | 0.058 | 0.010 | 0.007 | 0.007 | 0.004 | 2.0 | 0.173 | 43.1 | 12.4 | 37.9 | 10.6 | 2.1 | 0.116 |
| Heschl's Gyrus L | 0.64 | 0.07 | 0.66 | 0.06 | -1.3 | 0.389 | 0.80 | 0.09 | 0.74 | 0.09 | 3.1 | 0.018 | 0.006 | 0.005 | 0.007 | 0.004 | -1.1 | 0.526 | 30.6 | 12.9 | 33.9 | 11.2 | -1.3 | 0.379 |
| Temporal Fusiform Cortex; Anterior Division L | 0.54 | 0.06 | 0.57 | 0.11 | -1.3 | 0.396 | 0.56 | 0.25 | 0.59 | 0.16 | -0.5 | 0.711 | 0.003 | 0.003 | 0.003 | 0.003 | -1.1 | 0.527 | 13.8 | 9.6 | 18.5 | 10.6 | -2.2 | 0.107 |
| Temporal Fusiform Cortex; Posterior Division L | 0.58 | 0.06 | 0.59 | 0.10 | -0.6 | 0.790 | 0.69 | 0.14 | 0.64 | 0.13 | 1.8 | 0.146 | 0.004 | 0.003 | 0.005 | 0.004 | -1.3 | 0.459 | 19.9 | 9.4 | 22.7 | 10.0 | -1.3 | 0.356 |
| Temporal Occipital Fusiform Cortex L | 0.58 | 0.06 | 0.61 | 0.06 | -2.8 | 0.061 | 0.68 | 0.19 | 0.70 | 0.11 | -0.6 | 0.683 | 0.003 | 0.003 | 0.005 | 0.004 | -2.9 | 0.062 | 19.5 | 10.6 | 25.4 | 9.0 | -2.8 | 0.028 |
| Middle Temporal Gyrus; Temporooccipital Part L | 0.63 | 0.08 | 0.64 | 0.06 | -0.3 | 0.849 | 0.75 | 0.12 | 0.74 | 0.09 | 0.5 | 0.704 | 0.006 | 0.004 | 0.006 | 0.004 | 0.8 | 0.668 | 30.7 | 12.7 | 30.7 | 12.0 | 0.0 | 0.987 |
| Inferior Temporal Gyrus; Temporooccipital Part L | 0.60 | 0.07 | 0.61 | 0.05 | -0.9 | 0.599 | 0.69 | 0.19 | 0.70 | 0.10 | -0.5 | 0.735 | 0.004 | 0.003 | 0.005 | 0.004 | -1.2 | 0.480 | 24.2 | 12.0 | 25.7 | 9.3 | -0.6 | 0.752 |
| Occipital Fusiform Gyrus L | 0.64 | 0.09 | 0.66 | 0.07 | -1.4 | 0.344 | 0.74 | 0.12 | 0.72 | 0.12 | 0.6 | 0.688 | 0.006 | 0.005 | 0.007 | 0.005 | -0.7 | 0.734 | 30.7 | 16.6 | 34.7 | 13.3 | -1.3 | 0.377 |
| Supracalcarine Cortex L | 0.61 | 0.05 | 0.63 | 0.06 | -1.2 | 0.455 | 0.77 | 0.11 | 0.71 | 0.12 | 2.6 | 0.041 | 0.004 | 0.003 | 0.005 | 0.003 | -2.1 | 0.152 | 25.5 | 9.5 | 27.9 | 10.7 | -1.1 | 0.459 |
| Cuneal Cortex L | 0.61 | 0.07 | 0.62 | 0.06 | -0.7 | 0.726 | 0.74 | 0.15 | 0.71 | 0.13 | 1.0 | 0.496 | 0.004 | 0.004 | 0.005 | 0.003 | -0.4 | 0.843 | 25.2 | 13.5 | 26.8 | 11.0 | -0.6 | 0.753 |
| Lingual Gyrus L | 0.61 | 0.12 | 0.63 | 0.12 |  |  |  |  |  |  |  |  |  |  |  |  |  |  |  |  |  |  |  |  |

Table S3 continued

|  | Global Efficiency |  |  |  |  |  | Local Efficiency |  |  |  |  |  | Betweenness Centrality |  |  |  |  |  | Degree |  |  |  |  |  |
| --- | --- | --- | --- | --- | --- | --- | --- | --- | --- | --- | --- | --- | --- | --- | --- | --- | --- | --- | --- | --- | --- | --- | --- | --- |
|  | Younger |  | Older |  | Young vs Old |  | Younger |  | Older |  | Young vs Old |  | Younger |  | Older |  | Young vs Old |  | Younger |  | Older |  | Young vs Old |  |
|  | Mean | SD | Mean | SD | T | p_FDR | Mean | SD | Mean | SD | T | p_FDR | Mean | SD | Mean | SD | T | p_FDR | Mean | SD | Mean | SD | T | p_FDR |
| Frontal Orbital Cortex R | 0.72 | 0.07 | 0.69 | 0.06 | 1.7 | 0.243 | 0.78 | 0.07 | 0.75 | 0.08 | 2.2 | 0.081 | 0.011 | 0.006 | 0.009 | 0.005 | 1.7 | 0.264 | 46.6 | 14.0 | 41.5 | 11.7 | 1.9 | 0.182 |
| Frontal Pole R | 0.84 | 0.03 | 0.81 | 0.05 | 3.6 | 0.020 | 0.74 | 0.02 | 0.73 | 0.03 | 1.4 | 0.309 | 0.030 | 0.012 | 0.022 | 0.007 | 4.0 | 0.008 | 71.8 | 6.8 | 65.4 | 9.0 | 3.6 | 0.005 |
| Frontal Operculum Cortex R | 0.64 | 0.07 | 0.66 | 0.06 | -1.4 | 0.372 | 0.78 | 0.15 | 0.75 | 0.09 | 1.1 | 0.423 | 0.006 | 0.005 | 0.007 | 0.005 | -1.3 | 0.409 | 32.4 | 11.8 | 35.6 | 10.8 | -1.3 | 0.373 |
| Superior Frontal Gyrus R | 0.72 | 0.08 | 0.69 | 0.07 | 2.2 | 0.128 | 0.77 | 0.13 | 0.74 | 0.10 | 1.6 | 0.222 | 0.011 | 0.006 | 0.008 | 0.005 | 2.2 | 0.145 | 48.3 | 13.6 | 40.9 | 14.4 | 2.4 | 0.063 |
| Middle Frontal Gyrus R | 0.78 | 0.07 | 0.75 | 0.07 | 2.5 | 0.080 | 0.77 | 0.05 | 0.74 | 0.05 | 3.2 | 0.012 | 0.018 | 0.007 | 0.014 | 0.007 | 2.6 | 0.073 | 60.6 | 11.4 | 52.5 | 15.0 | 2.8 | 0.029 |
| Inferior Frontal Gyrus; Pars Triangularis R | 0.70 | 0.05 | 0.69 | 0.05 | 0.6 | 0.804 | 0.82 | 0.07 | 0.76 | 0.06 | 4.1 | 0.001 | 0.007 | 0.005 | 0.009 | 0.005 | -1.7 | 0.258 | 42.0 | 10.3 | 40.8 | 11.5 | 0.5 | 0.824 |
| Inferior Frontal Gyrus; Pars Opercularis R | 0.73 | 0.05 | 0.69 | 0.05 | 4.1 | 0.006 | 0.81 | 0.05 | 0.76 | 0.07 | 3.8 | 0.003 | 0.011 | 0.005 | 0.008 | 0.004 | 2.8 | 0.047 | 48.9 | 8.8 | 40.0 | 10.5 | 4.2 | 0.002 |
| Paracingulate Gyrus R | 0.72 | 0.06 | 0.68 | 0.06 | 3.5 | 0.018 | 0.80 | 0.05 | 0.77 | 0.06 | 2.3 | 0.063 | 0.010 | 0.006 | 0.008 | 0.005 | 2.3 | 0.112 | 48.1 | 12.1 | 38.8 | 11.3 | 3.6 | 0.005 |
| Insular Cortex R | 0.60 | 0.07 | 0.60 | 0.10 | 0.4 | 0.830 | 0.72 | 0.17 | 0.69 | 0.12 | 0.9 | 0.500 | 0.005 | 0.005 | 0.004 | 0.003 | 0.8 | 0.673 | 24.6 | 10.9 | 23.6 | 10.7 | 0.4 | 0.847 |
| Amygdala r | 0.56 | 0.12 | 0.58 | 0.07 | -1.1 | 0.528 | 0.58 | 0.28 | 0.58 | 0.18 | 0.0 | 0.992 | 0.005 | 0.006 | 0.005 | 0.004 | 0.2 | 0.908 | 20.1 | 14.4 | 19.6 | 10.3 | 0.2 | 0.940 |
| Juxtapositional Lobule Cortex R | 0.60 | 0.13 | 0.60 | 0.11 | -0.2 | 0.874 | 0.77 | 0.17 | 0.70 | 0.11 | 2.1 | 0.096 | 0.005 | 0.006 | 0.005 | 0.004 | 0.1 | 0.925 | 25.7 | 15.1 | 25.5 | 13.0 | 0.1 | 0.973 |
| Precentral Gyrus R | 0.73 | 0.06 | 0.70 | 0.08 | 1.9 | 0.185 | 0.79 | 0.05 | 0.73 | 0.07 | 4.5 | 0.000 | 0.013 | 0.007 | 0.011 | 0.007 | 1.0 | 0.553 | 48.6 | 12.2 | 42.9 | 15.9 | 1.8 | 0.184 |
| Postcentral Gyrus R | 0.66 | 0.07 | 0.66 | 0.08 | -0.2 | 0.870 | 0.80 | 0.10 | 0.68 | 0.17 | 3.7 | 0.004 | 0.006 | 0.004 | 0.008 | 0.005 | -1.9 | 0.202 | 34.2 | 13.0 | 35.3 | 16.6 | -0.3 | 0.857 |
| Central Opercular Cortex R | 0.62 | 0.10 | 0.65 | 0.06 | -1.9 | 0.184 | 0.74 | 0.15 | 0.72 | 0.10 | 0.5 | 0.694 | 0.006 | 0.007 | 0.007 | 0.004 | -0.4 | 0.850 | 28.4 | 14.6 | 33.1 | 11.3 | -1.7 | 0.244 |
| Superior Parietal Lobule R | 0.62 | 0.13 | 0.63 | 0.07 | -0.3 | 0.857 | 0.79 | 0.09 | 0.73 | 0.12 | 2.6 | 0.040 | 0.006 | 0.005 | 0.005 | 0.003 | 1.1 | 0.534 | 29.6 | 15.8 | 28.3 | 13.3 | 0.4 | 0.837 |
| Supramarginal Gyrus; Anterior Division R | 0.61 | 0.07 | 0.61 | 0.05 | 0.5 | 0.834 | 0.79 | 0.09 | 0.69 | 0.12 | 4.5 | 0.001 | 0.004 | 0.004 | 0.004 | 0.002 | 0.4 | 0.844 | 26.2 | 14.0 | 24.5 | 10.6 | 0.6 | 0.754 |
| Supramarginal Gyrus; Posterior Division R | 0.65 | 0.07 | 0.66 | 0.06 | -0.6 | 0.802 | 0.79 | 0.08 | 0.73 | 0.09 | 2.8 | 0.027 | 0.007 | 0.006 | 0.007 | 0.004 | 0.1 | 0.923 | 34.1 | 13.4 | 35.3 | 12.1 | -0.4 | 0.837 |
| Angular Gyrus R | 0.67 | 0.07 | 0.68 | 0.07 | -0.4 | 0.819 | 0.78 | 0.09 | 0.73 | 0.10 | 2.6 | 0.041 | 0.008 | 0.006 | 0.008 | 0.005 | 0.3 | 0.879 | 37.7 | 13.6 | 38.8 | 14.2 | -0.4 | 0.842 |
| Parietal Operculum Cortex R | 0.60 | 0.07 | 0.61 | 0.08 | -0.4 | 0.830 | 0.72 | 0.18 | 0.67 | 0.18 | 1.2 | 0.377 | 0.005 | 0.004 | 0.005 | 0.004 | -0.3 | 0.843 | 24.5 | 12.5 | 25.5 | 11.7 | -0.4 | 0.837 |
| Parahippocampal Gyrus; Anterior Division R | 0.54 | 0.08 | 0.57 | 0.10 | -1.5 | 0.299 | 0.50 | 0.23 | 0.57 | 0.18 | -1.7 | 0.188 | 0.005 | 0.005 | 0.004 | 0.002 | 1.5 | 0.347 | 14.5 | 8.9 | 18.4 | 8.3 | -2.1 | 0.116 |
| Parahippocampal Gyrus; Posterior Division R | 0.49 | 0.17 | 0.55 | 0.13 | -1.8 | 0.221 | 0.57 | 0.19 | 0.59 | 0.15 | -0.6 | 0.710 | 0.002 | 0.002 | 0.003 | 0.003 | -2.4 | 0.104 | 11.7 | 8.4 | 16.6 | 8.3 | -2.7 | 0.033 |
| Hippocampus R | 0.56 | 0.09 | 0.57 | 0.10 | -0.7 | 0.752 | 0.56 | 0.29 | 0.57 | 0.23 | -0.1 | 0.946 | 0.005 | 0.006 | 0.004 | 0.003 | 1.6 | 0.284 | 17.5 | 12.5 | 19.5 | 12.6 | -0.7 | 0.722 |
| Thalamus R | 0.55 | 0.11 | 0.59 | 0.11 | -1.7 | 0.242 | 0.68 | 0.18 | 0.65 | 0.19 | 0.8 | 0.607 | 0.003 | 0.003 | 0.004 | 0.005 | -2.0 | 0.177 | 15.8 | 11.5 | 22.4 | 13.4 | -2.4 | 0.062 |
| Caudate R | 0.56 | 0.07 | 0.61 | 0.11 | -2.7 | 0.062 | 0.61 | 0.23 | 0.68 | 0.15 | -1.6 | 0.213 | 0.003 | 0.003 | 0.005 | 0.003 | -3.1 | 0.062 | 16.6 | 10.4 | 27.7 | 11.5 | -4.7 | 0.001 |
| Putamen R | 0.61 | 0.09 | 0.66 | 0.12 | -2.0 | 0.173 | 0.67 | 0.16 | 0.73 | 0.06 | -2.2 | 0.083 | 0.009 | 0.008 | 0.009 | 0.006 | -0.3 | 0.848 | 26.9 | 15.8 | 36.3 | 14.3 | -2.9 | 0.022 |
| Pallidum R | 0.52 | 0.08 | 0.56 | 0.11 | -1.9 | 0.182 | 0.50 | 0.28 | 0.60 | 0.17 | -2.2 | 0.083 | 0.003 | 0.003 | 0.003 | 0.003 | -0.8 | 0.671 | 12.0 | 8.2 | 17.8 | 9.7 | -2.9 | 0.022 |
| Accumbens R | 0.53 | 0.07 | 0.57 | 0.10 | -2.3 | 0.124 | 0.53 | 0.23 | 0.63 | 0.12 | -2.5 | 0.050 | 0.002 | 0.002 | 0.003 | 0.002 | -1.9 | 0.206 | 11.9 | 6.7 | 18.3 | 8.6 | -3.8 | 0.005 |
| Temporal Pole R | 0.62 | 0.07 | 0.62 | 0.12 | 0.4 | 0.820 | 0.74 | 0.12 | 0.66 | 0.13 | 2.9 | 0.022 | 0.007 | 0.007 | 0.006 | 0.005 | 0.3 | 0.863 | 28.3 | 13.3 | 27.9 | 15.0 | 0.1 | 0.974 |
| Planum Polare R | 0.57 | 0.08 | 0.59 | 0.10 | -0.6 | 0.806 | 0.65 | 0.19 | 0.67 | 0.16 | -0.4 | 0.754 | 0.003 | 0.003 | 0.004 | 0.005 | -0.9 | 0.636 | 19.7 | 11.7 | 21.5 | 10.3 | -0.7 | 0.718 |
| Superior Temporal Gyrus; Anterior Division R | 0.55 | 0.07 | 0.59 | 0.10 | -2.0 | 0.160 | 0.65 | 0.22 | 0.70 | 0.12 | -1.4 | 0.282 | 0.002 | 0.002 | 0.004 | 0.002 | -2.8 | 0.044 | 15.9 | 10.2 | 22.7 | 9.4 | -3.3 | 0.012 |
| Superior Temporal Gyrus; Posterior Division R | 0.66 | 0.07 | 0.65 | 0.05 | 0.4 | 0.839 | 0.78 | 0.10 | 0.75 | 0.08 | 1.7 | 0.191 | 0.006 | 0.005 | 0.006 | 0.004 | 0.2 | 0.909 | 34.5 | 13.1 | 33.2 | 11.0 | 0.5 | 0.827 |
| Middle Temporal Gyrus; Anterior Division R | 0.64 | 0.06 | 0.63 | 0.05 | 1.1 | 0.521 | 0.73 | 0.15 | 0.73 | 0.08 | 0.0 | 0.988 | 0.008 | 0.006 | 0.005 | 0.003 | 2.5 | 0.079 | 31.1 | 12.3 | 28.0 | 8.4 | 1.4 | 0.338 |
| Middle Temporal Gyrus; Posterior Division R | 0.72 | 0.07 | 0.68 | 0.07 | 3.0 | 0.039 | 0.77 | 0.06 | 0.74 | 0.07 | 2.1 | 0.096 | 0.013 | 0.008 | 0.009 | 0.006 | 2.9 | 0.048 | 47.7 | 14.1 | 38.7 | 13.6 | 3.0 | 0.019 |
| Inferior Temporal Gyrus; Anterior Division R | 0.62 | 0.07 | 0.59 | 0.10 | 1.4 | 0.367 | 0.73 | 0.15 | 0.65 | 0.13 | 2.6 | 0.041 | 0.006 | 0.005 | 0.005 | 0.003 | 0.9 | 0.624 | 27.4 | 13.7 | 23.3 | 9.6 | 1.6 | 0.244 |
| Inferior Temporal Gyrus; Posterior Division R | 0.64 | 0.06 | 0.63 | 0.07 | 1.0 | 0.556 | 0.76 | 0.10 | 0.70 | 0.12 | 2.5 | 0.050 | 0.007 | 0.006 | 0.006 | 0.005 | 0.8 | 0.689 | 31.5 | 11.8 | 28.4 | 11.7 | 1.2 | 0.395 |
| Planum Temporale R | 0.67 | 0.06 | 0.67 | 0.05 | -0.2 | 0.886 | 0.80 | 0.07 | 0.75 | 0.08 | 3.0 | 0.022 | 0.008 | 0.005 | 0.008 | 0.005 | -0.2 | 0.873 | 37.1 | 12.0 | 37.4 | 9.7 | -0.1 | 0.965 |
| Heschl's Gyrus R | 0.63 | 0.07 | 0.65 | 0.06 | -1.6 | 0.272 | 0.75 | 0.12 | 0.75 | 0.09 | 0.0 | 1.000 | 0.006 | 0.005 | 0.006 | 0.004 | 0.0 | 0.982 | 28.4 | 13.1 | 32.5 | 9.5 | -1.7 | 0.241 |
| Temporal Fusiform Cortex; Anterior Division R | 0.53 | 0.08 | 0.57 | 0.10 | -2.0 | 0.178 | 0.55 | 0.23 | 0.59 | 0.16 | -1.0 | 0.473 | 0.003 | 0.004 | 0.004 | 0.003 | -0.7 | 0.691 | 12.5 | 8.2 | 18.1 | 9.4 | -2.9 | 0.022 |
| Temporal Fusiform Cortex; Posterior Division R | 0.58 | 0.07 | 0.59 | 0.10 | -0.5 | 0.836 | 0.62 | 0.18 | 0.62 | 0.13 | -0.1 | 0.989 | 0.006 | 0.007 | 0.005 | 0.003 | 0.5 | 0.799 | 20.0 | 10.8 | 22.3 | 9.6 | -1.0 | 0.487 |
| Temporal Occipital Fusiform Cortex R | 0.59 | 0.06 | 0.60 | 0.10 | -0.6 | 0.773 | 0.70 | 0.17 | 0.69 | 0.12 | 0.5 | 0.733 | 0.005 | 0.006 | 0.005 | 0.004 | 0.0 | 0.981 | 21.8 | 11.0 | 24.8 | 9.9 | -1.3 | 0.365 |
| Middle Temporal Gyrus; Temporooccipital Part R | 0.65 | 0.06 | 0.65 | 0.05 | -0.1 | 0.910 | 0.75 | 0.11 | 0.73 | 0.08 | 0.9 | 0.522 | 0.007 | 0.005 | 0.007 | 0.004 | 0.4 | 0.840 | 33.1 | 13.1 | 33.3 | 11.1 | -0.1 | 0.966 |
| Inferior Temporal Gyrus; Temporooccipital Part R | 0.60 | 0.06 | 0.61 | 0.06 | -0.5 | 0.813 | 0.71 | 0.14 | 0.69 | 0.14 | 0.9 | 0.516 | 0.006 | 0.006 | 0.005 | 0.003 | 0.9 | 0.632 | 24.2 | 11.4 | 25.2 | 10.0 | -0.4 | 0.841 |
| Occipital Fusiform Gyrus R | 0.64 | 0.08 | 0.65 | 0.06 | -0.8 | 0.706 | 0.73 | 0.11 | 0.72 | 0.11 | 0.5 | 0.698 | 0.007 | 0.007 | 0.007 | 0.005 | 0.5 | 0.825 | 31.4 | 14.8 | 33.3 | 12.7 | -0.7 | 0.761 |
| Supracalcarine Cortex R | 0.65 | 0.07 | 0.63 | 0.05 | 1.3 | 0.399 | 0.74 | 0.14 | 0.72 | 0.10 | 0.7 | 0.642 | 0.006 | 0.004 | 0.005 | 0.004 | 0.8 | 0.666 | 32.8 | 13.3 | 28.7 | 9.9 | 1.6 | 0.248 |
| Cuneal Cortex R | 0.63 | 0.06 | 0.64 | 0.06 | -0.4 | 0.827 | 0.78 | 0.10 | 0.72 | 0.12 | 2.2 | 0.083 | 0.005 | 0.004 | 0.006 | 0.004 | -0.8 | 0.659 | 29.8 | 12.8 | 30.6 | 10.7 | -0.3 | 0.864 |
| Lingual Gyrus R | 0.60 | 0.12 | 0.63 | 0.11 | -0.9 | 0.597 | 0.72 | 0.13 | 0.69 | 0.13 | 1.3 | 0.338 | 0.005 | 0.004 |  |  |  |  |  |  |  |  |  |  |

### 2.5 Age Differences in the Glucose Cost Index

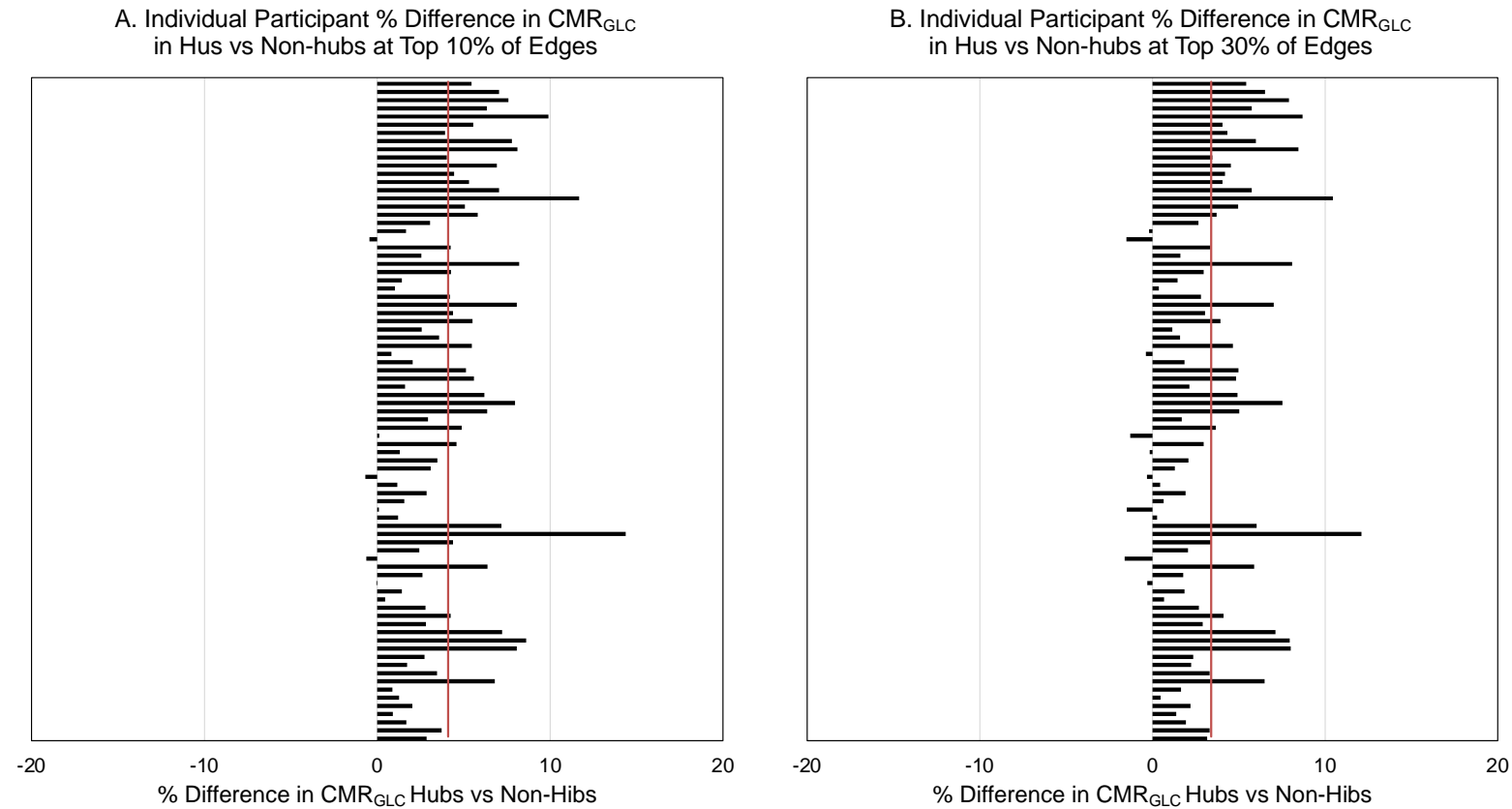

**Figure S5.** Individual participant percentage difference in cerebral metabolic rates of glucose in hub and non-hub regions at the (A) top 10% and (B) top 30% of edges. Red vertical lines are the sample average. The average $CMR_{GLC}$ was 4.1% higher in the 13 hubs than the 93 non-hub regions at the top 10% of edges ($t(80) = 12.7$, $p < .001$). The average $CMR_{GLC}$ was also higher in the hub than non-hub regions for 83 of 86 participants. The average $CMR_{GLC}$ was 3.4% higher in the 15 hubs than the 91 non-hub regions at the top 30% of edges ($t(80) = 10.8$, $p < .001$). It was also in the hub vs non-hub regions higher for 78 of 86 participants.

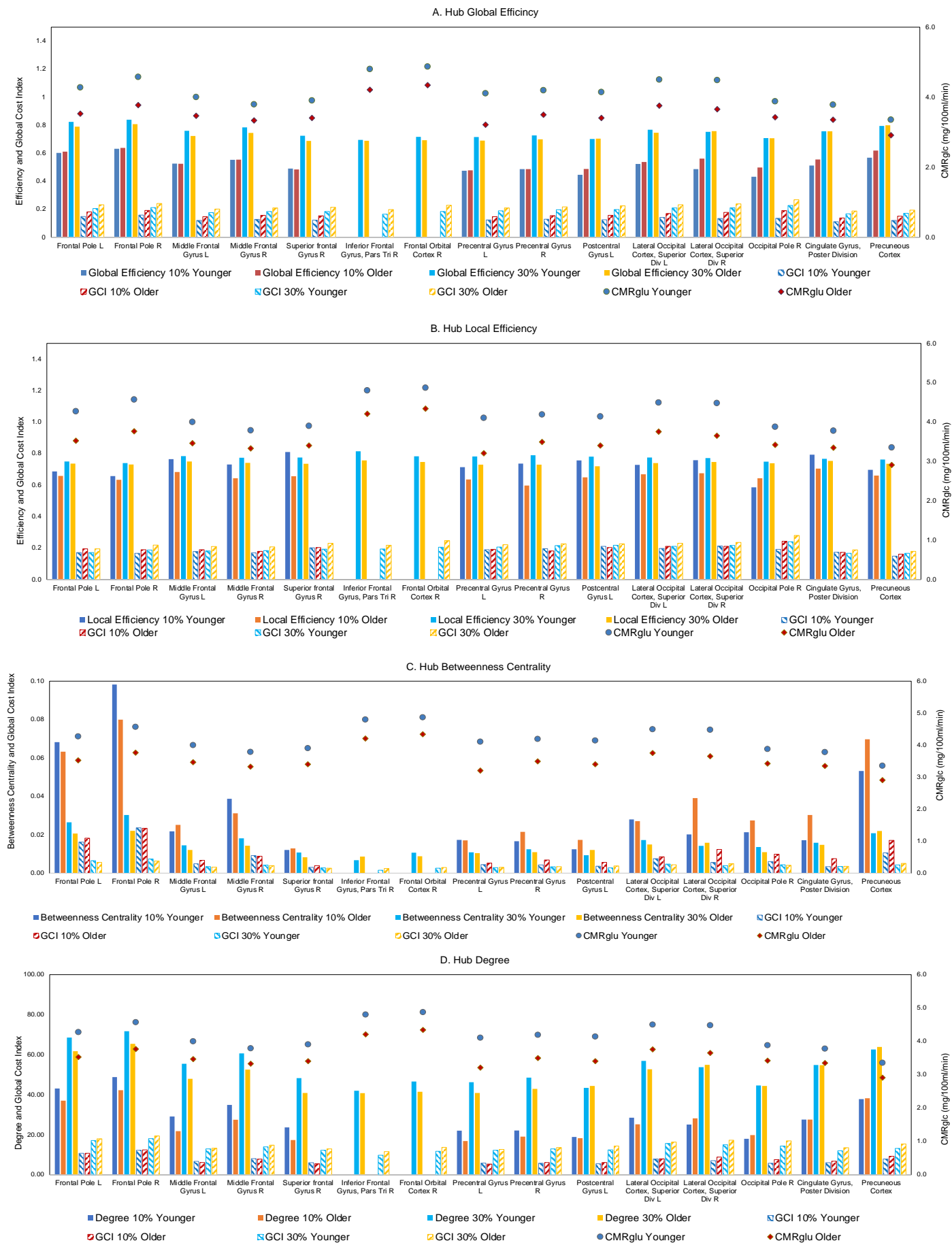

**Figure S6.** CMR<sub>GLU</sub> indexed across the entire scan, and mean graph metrics and glucose cost index (GCI) in hub regions for younger and older adults. (A) Global efficiency, (B) local efficiency, (C), betweenness centrality, and (D) degree are shown at top 10% and top 30% of edges. The significant age group differences on the graph metrics and glucose cost index are plotted on glass brains in Figure 2 and Figure 3, respectively, in the main manuscript.

**Table S4.** Mean and Standard Deviation (SD) and GLM results of average Glucose Cost Index (GCI) for each graph metric at top 10% and 30% of edges, and results of GLMs comparing CGI at 10% and 30% of edge threshold and hubs vs non-hubs.

|  | 10% Edges |  |  |  | 30% Edges |  |  |  | F-Test |  |  |  |
| --- | --- | --- | --- | --- | --- | --- | --- | --- | --- | --- | --- | --- |
|  | Hubs |  | Non-Hubs |  | Hubs |  | Non-Hubs |  | 10% vs 30% Edges |  | Hubs vs Non-Hubs |  |
|  | Mean | SD | Mean | SD | Mean | SD | Mean | SD | F | p | F | p |
| GCI: Global Efficiency | 0.53 | 0.05 | 0.36 | 0.08 | 0.74 | 0.03 | 0.62 | 0.04 | 849.9 | 0.000 | 1401.3 | 0.000 |
| GCI: Local Efficiency | 0.68 | 0.09 | 0.56 | 0.16 | 0.75 | 0.04 | 0.70 | 0.04 | 155.8 | 0.000 | 127.4 | 0.000 |
| GCI: Betweenness Centrality | 0.034 | 0.010 | 0.008 | 0.003 | 0.015 | 0.003 | 0.006 | 0.001 | 1114.7 | 0.000 | 250.3 | 0.000 |
| GCI: Degree | 27.1 | 4.4 | 8.2 | 0.6 | 51.2 | 5.8 | 28.2 | 1.0 | 1133.6 | 0.000 | 45764.5 | 0.000 |

**Table S5.** Mean, Standard Deviation (SD) and post-hoc univariate F-test (df = 1,84) of age group differences in glucose efficiency index in the metabolic hubs at top 10% and 30% of network edges. F-values are plotted on the glass brains in Figures 3 in the main manuscript.

|  | Glucose Cost Index: Global Efficiency |  |  |  |  |  | Glucose Cost Index: Local Efficiency |  |  |  |  |  | Glucose Cost Index: Betweenness Centrality |  |  |  |  |  | Glucose Cost Index: Degree |  |  |  |  |  |
| --- | --- | --- | --- | --- | --- | --- | --- | --- | --- | --- | --- | --- | --- | --- | --- | --- | --- | --- | --- | --- | --- | --- | --- | --- |
|  | Younger |  | Older |  | Younger Vs Older |  | Younger |  | Older |  | Younger Vs Older |  | Younger |  | Older |  | Younger Vs Older |  | Younger |  | Older |  | Younger Vs Older |  |
|  | Mean | SD | Mean | SD | F | p | Mean | SD | Mean | SD | F | p | Mean | SD | Mean | SD | F | p | Mean | SD | Mean | SD | F | p |
| Top 10% of Edges |  |  |  |  |  |  |  |  |  |  |  |  |  |  |  |  |  |  |  |  |  |  |  |  |
| Frontal Pole L | 0.15 | 0.04 | 0.18 | 0.04 | 14.1 | 0.000 | 0.17 | 0.04 | 0.20 | 0.05 | 9.7 | 0.003 | 0.016 | 0.009 | 0.018 | 0.010 | 0.9 | 0.352 | 10.7 | 3.7 | 10.7 | 3.5 | 0.0 | 0.982 |
| Frontal Pole R | 0.16 | 0.04 | 0.19 | 0.04 | 12.5 | 0.000 | 0.16 | 0.03 | 0.19 | 0.03 | 12.1 | 0.001 | 0.024 | 0.014 | 0.023 | 0.012 | 0.0 | 0.914 | 12.3 | 3.9 | 12.5 | 3.9 | 0.0 | 0.834 |
| Middle Frontal Gyrus L | 0.12 | 0.03 | 0.15 | 0.03 | 15.6 | 0.000 | 0.17 | 0.04 | 0.19 | 0.06 | 2.7 | 0.104 | 0.005 | 0.006 | 0.007 | 0.006 | 2.0 | 0.165 | 6.7 | 2.8 | 6.1 | 2.8 | 0.9 | 0.347 |
| Middle Frontal Gyrus R | 0.13 | 0.04 | 0.16 | 0.04 | 11.9 | 0.000 | 0.16 | 0.04 | 0.18 | 0.05 | 2.8 | 0.097 | 0.009 | 0.008 | 0.009 | 0.007 | 0.0 | 0.863 | 8.1 | 2.7 | 7.8 | 3.7 | 0.2 | 0.685 |
| Superior frontal Gyrus R | 0.12 | 0.04 | 0.15 | 0.04 | 10.9 | 0.000 | 0.20 | 0.04 | 0.20 | 0.10 | 0.0 | 0.914 | 0.003 | 0.003 | 0.004 | 0.004 | 1.8 | 0.188 | 6.0 | 3.2 | 5.5 | 3.4 | 0.5 | 0.462 |
| Precentral Gyrus L | 0.12 | 0.04 | 0.15 | 0.05 | 5.6 | 0.000 | 0.18 | 0.07 | 0.19 | 0.09 | 0.5 | 0.471 | 0.005 | 0.004 | 0.005 | 0.006 | 0.6 | 0.457 | 5.8 | 3.5 | 5.4 | 4.1 | 0.3 | 0.585 |
| Precentral Gyrus R | 0.13 | 0.03 | 0.15 | 0.04 | 8.3 | 0.000 | 0.20 | 0.06 | 0.18 | 0.08 | 1.1 | 0.304 | 0.004 | 0.004 | 0.007 | 0.007 | 4.4 | 0.040 | 5.8 | 3.0 | 6.1 | 4.3 | 0.2 | 0.671 |
| Postcentral Gyrus L | 0.12 | 0.05 | 0.16 | 0.05 | 8.4 | 0.000 | 0.20 | 0.07 | 0.20 | 0.08 | 0.0 | 0.989 | 0.004 | 0.005 | 0.006 | 0.005 | 2.9 | 0.091 | 5.5 | 4.0 | 6.1 | 4.0 | 0.5 | 0.500 |
| Lateral Occipital Cortex, Superior Division L | 0.14 | 0.04 | 0.17 | 0.04 | 8.9 | 0.000 | 0.19 | 0.04 | 0.21 | 0.06 | 0.6 | 0.433 | 0.008 | 0.008 | 0.009 | 0.006 | 0.4 | 0.546 | 7.9 | 3.9 | 8.0 | 3.7 | 0.0 | 0.926 |
| Lateral Occipital Cortex, Superior Division R | 0.13 | 0.05 | 0.18 | 0.04 | 17.6 | 0.000 | 0.20 | 0.05 | 0.21 | 0.04 | 0.2 | 0.639 | 0.006 | 0.006 | 0.012 | 0.009 | 15.8 | 0.000 | 7.1 | 4.0 | 8.8 | 4.1 | 3.8 | 0.055 |
| Occipital Pole R | 0.13 | 0.06 | 0.19 | 0.07 | 14.8 | 0.000 | 0.19 | 0.10 | 0.24 | 0.09 | 5.0 | 0.029 | 0.006 | 0.007 | 0.010 | 0.010 | 3.7 | 0.057 | 5.8 | 4.9 | 7.6 | 5.7 | 2.3 | 0.137 |
| Cingulate Gyrus, Poster Division | 0.11 | 0.04 | 0.14 | 0.04 | 10.0 | 0.000 | 0.17 | 0.04 | 0.17 | 0.03 | 0.2 | 0.620 | 0.003 | 0.004 | 0.008 | 0.005 | 16.3 | 0.000 | 6.0 | 2.6 | 6.8 | 2.8 | 1.8 | 0.179 |
| Precuneous Cortex | 0.12 | 0.04 | 0.15 | 0.03 | 16.5 | 0.000 | 0.15 | 0.03 | 0.16 | 0.03 | 1.5 | 0.233 | 0.011 | 0.008 | 0.017 | 0.011 | 9.5 | 0.003 | 7.9 | 2.9 | 9.3 | 2.9 | 4.8 | 0.032 |
| Top 30% of Edges |  |  |  |  |  |  |  |  |  |  |  |  |  |  |  |  |  |  |  |  |  |  |  |  |
| Frontal Pole L | 0.20 | 0.06 | 0.23 | 0.04 | 6.5 | 0.012 | 0.17 | 0.05 | 0.20 | 0.05 | 5.3 | 0.025 | 0.007 | 0.004 | 0.006 | 0.002 | 1.7 | 0.200 | 17.1 | 5.5 | 18.0 | 4.0 | 0.7 | 0.414 |
| Frontal Pole R | 0.21 | 0.05 | 0.24 | 0.04 | 6.9 | 0.010 | 0.19 | 0.05 | 0.22 | 0.04 | 9.1 | 0.004 | 0.007 | 0.003 | 0.006 | 0.002 | 3.6 | 0.061 | 18.1 | 5.0 | 19.4 | 4.4 | 1.7 | 0.201 |
| Middle Frontal Gyrus L | 0.18 | 0.04 | 0.20 | 0.04 | 8.0 | 0.006 | 0.18 | 0.05 | 0.21 | 0.05 | 6.9 | 0.010 | 0.003 | 0.002 | 0.003 | 0.001 | 0.9 | 0.355 | 12.9 | 3.4 | 13.3 | 4.0 | 0.2 | 0.643 |
| Middle Frontal Gyrus R | 0.18 | 0.04 | 0.21 | 0.04 | 7.3 | 0.008 | 0.18 | 0.06 | 0.21 | 0.04 | 4.6 | 0.036 | 0.004 | 0.002 | 0.004 | 0.002 | 0.2 | 0.637 | 14.0 | 4.0 | 14.8 | 5.1 | 0.6 | 0.426 |
| Superior frontal Gyrus R | 0.18 | 0.05 | 0.21 | 0.05 | 8.7 | 0.004 | 0.19 | 0.06 | 0.23 | 0.05 | 8.9 | 0.004 | 0.003 | 0.002 | 0.003 | 0.001 | 0.3 | 0.557 | 12.2 | 4.5 | 13.0 | 5.1 | 0.5 | 0.494 |
| Inferior Frontal Gyrus, Pars Triangular R | 0.16 | 0.04 | 0.20 | 0.04 | 14.0 | 0.000 | 0.19 | 0.06 | 0.22 | 0.05 | 4.2 | 0.044 | 0.002 | 0.001 | 0.002 | 0.001 | 9.7 | 0.003 | 9.8 | 3.5 | 11.6 | 4.0 | 4.4 | 0.039 |
| Frontal Orbital Cortex R | 0.18 | 0.04 | 0.23 | 0.05 | 16.3 | 0.000 | 0.21 | 0.06 | 0.25 | 0.07 | 7.4 | 0.008 | 0.003 | 0.002 | 0.003 | 0.002 | 0.6 | 0.457 | 11.8 | 4.0 | 13.7 | 5.5 | 3.1 | 0.082 |
| Precentral Gyrus L | 0.19 | 0.05 | 0.21 | 0.05 | 3.4 | 0.068 | 0.21 | 0.06 | 0.22 | 0.05 | 1.2 | 0.276 | 0.003 | 0.002 | 0.003 | 0.002 | 0.1 | 0.719 | 12.3 | 5.6 | 12.7 | 5.8 | 0.1 | 0.757 |
| Precentral Gyrus R | 0.20 | 0.05 | 0.22 | 0.05 | 3.7 | 0.059 | 0.22 | 0.06 | 0.23 | 0.05 | 0.8 | 0.386 | 0.003 | 0.002 | 0.003 | 0.002 | 0.1 | 0.797 | 12.9 | 3.8 | 13.6 | 5.9 | 0.4 | 0.514 |
| Postcentral Gyrus L | 0.20 | 0.06 | 0.22 | 0.05 | 4.6 | 0.035 | 0.22 | 0.06 | 0.23 | 0.06 | 0.4 | 0.547 | 0.003 | 0.002 | 0.004 | 0.002 | 4.3 | 0.041 | 12.5 | 6.2 | 14.4 | 6.3 | 1.9 | 0.177 |
| Lateral Occipital Cortex, Superior Division L | 0.21 | 0.06 | 0.23 | 0.05 | 3.4 | 0.067 | 0.21 | 0.06 | 0.23 | 0.05 | 2.6 | 0.114 | 0.005 | 0.003 | 0.004 | 0.002 | 0.4 | 0.513 | 15.6 | 5.7 | 16.4 | 5.0 | 0.4 | 0.546 |
| Lateral Occipital Cortex, Superior Division R | 0.21 | 0.06 | 0.24 | 0.05 | 5.5 | 0.022 | 0.22 | 0.06 | 0.24 | 0.05 | 2.0 | 0.159 | 0.004 | 0.002 | 0.005 | 0.003 | 3.8 | 0.056 | 15.0 | 5.7 | 17.3 | 5.3 | 3.6 | 0.062 |
| Occipital Pole R | 0.23 | 0.07 | 0.27 | 0.08 | 6.1 | 0.016 | 0.24 | 0.07 | 0.28 | 0.09 | 4.5 | 0.037 | 0.004 | 0.004 | 0.004 | 0.003 | 0.2 | 0.673 | 14.4 | 7.0 | 16.9 | 8.1 | 2.3 | 0.137 |
| Cingulate Gyrus, Poster Division | 0.17 | 0.04 | 0.19 | 0.04 | 5.8 | 0.018 | 0.17 | 0.05 | 0.19 | 0.05 | 3.9 | 0.052 | 0.003 | 0.003 | 0.004 | 0.002 | 0.1 | 0.759 | 12.0 | 4.4 | 13.5 | 3.4 | 3.1 | 0.081 |
| Precuneous Cortex | 0.17 | 0.04 | 0.19 | 0.04 | 7.5 | 0.008 | 0.17 | 0.05 | 0.18 | 0.04 | 1.5 | 0.220 | 0.004 | 0.002 | 0.005 | 0.002 | 3.0 | 0.089 | 13.2 | 3.5 | 15.4 | 3.7 | 7.7 | 0.007 |

Age group differences were found in the multivariate test at the top 10% of edges for global efficiency ($F(15,70) = 2.7$, $p = .004$), local efficiency ($F(15,70) = 3.4$, $p = .001$), betweenness centrality ($F(15,70) = 3.2$, $p = .001$) but not degree ($F(15,70) = 1.5$, $p = .158$). Age differences were also found at the top 30% of edges for global efficiency ($F(15,70) = 2.0$, $p = .002$), local efficiency ($F(15,70) = 2.7$, $p = .003$), betweenness centrality, ($F(15,70) = 3.5$, $p < .001$) but not degree ($F(15,70) = 1.4$, $p = .176$).
